# Hierarchical microstructural tissue growth of the gray and white matter of human visual cortex during the first year of life

**DOI:** 10.1101/2025.06.20.660718

**Authors:** Karla Perez, Ahmad Allen, Christina Tyagi, Sarah S. Tung, Bella Fascendini, Xiaoqian Yan, Juliet Horenziak, Danya Ortiz, Hua Wu, Kalanit Grill-Spector, Vaidehi S. Natu

**Affiliations:** Department of Psychology, Stanford University, Stanford, CA, 94305, USA; Department of Psychology, Princeton University, Princeton, NJ, 08544, USA; Institute of Science and Technology for Brain-Inspired Intelligence, Fudan University, Shanghai, China; Center for Cognitive and Neurobiological Imaging, Stanford, CA, 94305, USA; Wu Tsai Neurosciences Institute, Stanford University, Stanford, CA, 94305, USA; Neurosciences Program, Stanford University, Stanford, CA, 94305, USA

**Keywords:** development, gray-white matter, infants, microstructure, visual streams

## Abstract

Development of gray and white matter tissue microstructure is critical for the emergence of sensory and cognitive functions. However, it is unknown how microstructural tissue properties of the human visual system develop in the first year of human life. Here, we use tissue relaxation rate (R_1_) obtained using quantitative MRI to measure the longitudinal development of gray and white matter in brain areas spanning three visual processing streams: dorsal, lateral, and ventral, during the first year of life. R_1_ in gray and white matter of all visual regions in the three processing streams increases postnatally, indicating microstructural tissue growth. R_1_ increases faster between 0-6 months than 6-12 months, and faster in white matter than gray matter, with white matter R_1_ surpassing that of gray matter after two months of age. Strikingly, this microstructural growth is hierarchical: across all streams, early visual areas are more mature at birth than higher-level areas but develop more slowly postnatally than higher-level areas. The exception is TO1 (MT) which is similar to V1: it is microstructurally more mature at birth and develops slower than neighboring areas. Overall, our findings provide the first comprehensive measurement of microstructural tissue growth in infancy across three visual processing streams and propose a new hypothesis that functional development of the visual cortex may be coupled with microstructural development and follows a similar hierarchical trajectory.

## Introduction

The first year of an infant’s life is a critical period of brain development. It is marked by rapid and dynamic changes in brain anatomy(Bethlehem et al. 2022; Tung et al. 2025), cortical microstructure(Ball et al. 2013; Zomeno et al. 2016; Lebenberg et al. 2019; Natu et al. 2021b), fiber connections(Kinney et al. 1988; Paus et al. 2001; Dubois et al. 2014; Grotheer et al. 2022; Kubota et al. 2025), and sensory-motor functions(Norcia and Tyler 1985; Norcia et al. 1990; Fausey et al. 2016; Smith et al. 2018; Ellis et al. 2021). In the visual domain, by six months of age, infants perceive color(Maule et al. 2023) and motion aspects of stimuli like direction and speed(Agyei et al. 2016), develop spatial and contrast sensitivities(Norcia et al. 1990; Morrone et al. 1990), and show a preference for face-like over non-face-like stimuli(Fantz 1963; Cassia et al. 2004). The normative development of such myriad and complex visual functions is thought to be associated with microstructural tissue development of visual areas supporting these functions. In gray matter, microstructure includes neurons, dendritic and synaptic processes(Huttenlocher et al. 1982; Katz and Shatz 1996; Elston et al. 2009; Elston and Fujita 2014), and myelin(Miller et al. 2012; Natu et al. 2019), whereas in white matter microstructure predominantly includes axonal sheaths and the myelin wrapped around them(Sherman and Brophy 2005). However, it is unknown what is the rate and sequence of microstructural development of the gray and white matter of visual areas supporting these developing visual abilities during the first year of human life.

Here, we will fill this gap in knowledge by investigating the microstructural development of gray and adjacent white matter, cross-sectionally and longitudinally, in visual areas spanning three processing streams – ventral, dorsal, and lateral visual processing streams. The dorsal visual stream projects from early visual areas to parietal cortex and is involved in spatial vision, visually guided action, and attention(Ungerleider and Mishkin 1982; Goodale et al. 1991; Rizzolatti and Matelli 2003). The lateral stream projects from early visual areas to lateral occipital temporal cortex (LOTC) and superior temporal cortex and is associated with visual dynamics, action and social understanding, and multimodal processing(Weiner and Grill-Spector 2013; Pitcher and Ungerleider 2021; Wurm and Caramazza 2022). The ventral stream extends anatomically from early visual areas (V1-V3) to ventral (inferior) occipital and temporal cortex, and is involved in perception, object recognition and categorization(Ungerleider and Mishkin 1982; Ungerleider and Haxby 1994; Milner and Goodale 2008; Wurm and Caramazza 2022). To measure microstructural development in vivo we utilize quantitative magnetic resonance imaging (qMRI)(Mezer et al. 2013; Lutti et al. 2014; Edwards et al. 2018; Weiskopf et al. 2021) of longitudinal relaxation rate (R_1_[s^−1^]). We measure R_1_ as in our prior infant research(Natu et al. 2021b; Grotheer et al. 2022; Tung et al. 2025) because R_1_ is directly proportional to microstructural tissue density: higher R_1_ is coupled with higher density. Thus, the R_1_ metric can inform if during development, gray and white matter become microstructurally dense.

### How does brain microstructure develop?

Prior histological research in post-mortem human and primate brain tissue found synaptogenesis and growth immediately after birth (Katz and Shatz 1996; Elston and Fujita 2014) followed by synaptic pruning and elimination of weak and redundant connections(Huttenlocher and Dabholkar 1997). These findings predict early microstructural growth followed by pruning and reduction in tissue, that is, an initial increase in R_1_ followed by a reduction in R_1_. The second hypothesis predicts tissue growth(Deoni et al. 2011; Ball et al. 2013; Natu et al. 2021b; Grotheer et al. 2022) related to formation of new synapses and dendritic arborization(Katz and Shatz 1996; Elston and Fujita 2014) as well as myelination(Miller et al. 2012; Natu et al. 2019, 2021b), predicting developmental increases in R_1_.

### Is microstructural development uniform across the visual system or heterogeneous?

One possibility is that the entire visual system will develop together in infancy as visual experience only begins after birth and may drive the microstructural development of the entire visual system. Alternatively, there may be heterogeneous development across visual processing hierarchies or across streams as different visual behaviors develop at different stages and rates during infancy and childhood. For example, functions associated with early visual cortex (e.g., contrast, spatial frequency)(Norcia and Tyler 1985; Norcia et al. 1990) mature earlier in infancy than high-level functions like recognition or spatial attention. Furthermore, there is evidence for earlier development of microstructure and fiber tracts in the ventral visual stream than the dorsal stream during childhood (ages 5-8)(Loenneker et al. 2011; Vinci-Booher et al. 2022), and it is possible this differential development emerges in infancy.

### Does the development of gray and adjacent, superficial, white matter of the visual system occur in tandem or does each tissue type have its own developmental trajectory during the first year of life?

Prior work suggests increase in myelination mainly drives white matter development myelination(Dubois et al. 2014; Fields 2015; Turner 2019; Grotheer et al. 2022), but gray matter development is influenced by multiple mechanisms like myelination, synaptic arborization, and dendritic proliferation. Hence, we predict different mechanisms in gray versus white matter of brain areas will have varying impacts on the microstructural tissue measure R_1_ reflected in its differential developmental rates during infancy.

To examine these hypotheses, we collected, during natural sleep, longitudinal and cross-sectional qMRI data in full-term, healthy infants that were between newborn and approximately one year of age. We examined microstructural development of the gray matter and superficial white matter of 18 retinotopic areas spanning the three visual processing streams. We assessed the development of R_1_ in the gray and superficial white matter per area and compared R_1_ development across areas and tissue types.

## Methods and Materials

### Participants

Eighty-two full term and healthy infants (N_female_ = 35) were enrolled as participants in the study. Data from infants who were unable to fall asleep in the MRI scanner were excluded due to excessive movement that compromised data quality. Forty-seven out of the eighty-two infants provided usable data. Therefore, we analyzed longitudinal and cross-sectional data from 47 infants (N_female_ = 20) across 85 sessions. 25 out of 47 infants were longitudinal, participating in two or more timepoints (**Supplementary Table 1** and **Supplementary Fig. 1**). Acquisition spanned four timepoints: newborns (*N_sessions_* = 27; M_age_ ± SD_age_: 29.14 ± 9.92 days), 3-month-olds (*N_sessions_* = 23; 106.69 ± 19.01 days), 6-month-olds (*N_sessions_* = 23; 189.56 ± 15.90 days), and 1-year-olds (*N_sessions_*= 12; 387.58 ± 36.78 days). The study included a diverse cohort of infants: 2 Hispanic, 6 Asian, 24 White, and 15 multiracial participants.

### Expectant parent and infant screening procedure

Expectant parents and their infants were recruited from the San Francisco Bay Area through social media outreach. A two-phase screening procedure was implemented. Initially, parents were interviewed via phone to determine eligibility based on exclusion criteria aimed at enrolling typically developing infants. Those who met the criteria were rescreened after childbirth. Mothers were excluded if they reported recreational drug use during pregnancy, substantial alcohol consumption (defined as more than three instances of alcohol consumption per trimester; more than 1 drink per occasion), a lifelong diagnosis of autism spectrum disorder or any condition involving psychosis or mania, or if they were on medications for such conditions during pregnancy. Additional exclusions included limited proficiency in written and spoken English to understand the instructions of the study or any learning disabilities that might interfere with study participation. Infants were excluded if they were born preterm (<37 gestational weeks) or low birthweight (<5 lbs 8 oz). Additional infant exclusion criteria included the presence of congenital, genetic, or neurological disorders, visual impairments, complications at birth requiring intensive care (e.g., NICU admission), prior head injuries, or any contraindications for MRI (e.g., metal implants). The study was approved by the Stanford University Institutional Review Board for Human Subjects Research. Participants received 30 dollars per hour as compensation for their participation.

### Data Acquisition

All infants participated in a series of scanning procedures using a 3T GE MRI system at the Center for Cognitive and Neurobiological Imaging, Stanford University, to acquire both anatomical and quantitative data. To prioritize infant safety, all scans were conducted under first-level SAR conditions. Of the 85 scanning sessions, 69 were collected using a Nova 32-channel head coil, and the remaining used a custom 32-channel infant head coil(Ghotra et al. 2021). MRI scanning sessions were held in the evenings, timed to align with each infant’s typical bedtime. Each session spanned approximately 2.5-5 hours, which included time for preparation and for the infant to fall asleep. Upon arrival, caregivers provided written informed consent for both themselves and their infant. Prior to entering the scanner, both the caregiver and infant were screened for metal, and the infants were dressed in MRI-compatible cotton onesies and footed pants supplied by the research team.

To minimize movement, infants were securely swaddled with their arms by their sides to prevent loop formation, and soft wax earplugs were placed in their ears. For sessions involving newborns, a MR-safe plastic immobilizer (MedVac, www.supertechx-ray.com) was used to help stabilize both body and head position. Once the infant was settled, the caregiver and infant entered the MRI suite together. Caregivers were encouraged to follow their child’s typical sleep routine. After the infant fell asleep, they were positioned on the scanner bed by the caregiver. Weighted bags were placed along the bed’s edges to reduce side-to-side movement, and additional padding was used around the head and body for further stabilization. MRI-safe neonatal noise attenuator (https://natus.com/) and infant headphones (https://www.alpinehearingprotection.com/products/muffy-baby) were placed over the ears to protect the infants’ hearing. An experimenter remained inside the MRI suite throughout the scan to monitor the infant. An infrared camera was mounted on the head coil to monitor the infant’s movements and head position in real-time. The MRI operator watched the live video feed and paused the scan if any signs of discomfort or waking were observed. Head motion was closely tracked, and scans were repeated if movement exceeded acceptable thresholds. In addition, each image sequence was assessed after acquisition, and repeated when necessary to ensure high-quality data.

### Data acquisition parameters and preprocessing

#### Anatomical MRI

T1-weighted and T2-weighted images were acquired and used for tissue segmentation. T1-weighted image acquisition parameters: TE = 122.77 ms; TR = 3650 ms; echo train length = 1; voxel size = 1 mm^3^; Scan time: ∼3 min. T2-weighted image acquisition parameters: TE =122 ms; TR = 3650 ms; echo train length = 120; voxel size = 1 mm^3^; FOV = 20.5 cm; Scan time: ∼4 min.

#### Quantitative MRI

An inversion-recovery EPI (IR-EPI) sequence with multiple inversions times (TI) was used to estimate quantitative relaxation time R_1_ (R_1_ = 1/T_1_) in each voxel. The IR-EPI used a slice-shuffling technique: 20 TIs with the first TI = 50 ms and TI interval = 150 ms, and a second IR-EPI with reverse-phase encoding direction. Other acquisition parameters were voxel size = 2 mm^3^; number of slices = 60; FOV = 20 cm; in-plane/through-plane acceleration = 1/3; Scan time: 1 min and 45 sec. To obtain R_1_ maps, we performed susceptibility-induced distortion correction on the IR-EPI images using FSL’s top-up and the IR-EPI acquisition with reverse-phase encoding direction. We then used the distortion corrected images to fit the T_1_ relaxation signal model using a multi-dimensional Levenberg-Marquardt algorithm. In an inversion-recovery sequence, the signal *S (t)* has an exponential decay over time (*t*) with a decay constant *T_1_*:

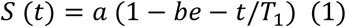

In Eq 1, *a* is a constant that is proportional to the initial magnetization of the voxel and *be* is the effective inversion coefficient of the voxel (for perfect inversion *b* = 2). We applied an absolute value operation on both sides of the equation and used the resulting equation as the fitting model. We use the absolute value of the signal equation because we apply the magnitude images to fit the model. The magnitude images only keep the information about the strength of the signal but not the phase or the sign of the signal. The output of the algorithm is the estimated quantitative T_1_ value per voxel. From the T_1_ estimate, we calculate R_1_ (R_1_ = 1/T_1_) as R_1_ is directly proportional to macromolecular tissue volume.

### MRI data preprocessing pipeline

#### Generation of cortical surfaces

We generated gray and white matter tissue segmentations using the T1- and T2-weighted images. Multiple steps were applied to generate an accurate segmentation of each infant’s brain at each timepoint (**Fig. 1a**): (1) An initial segmentation of gray and white matter was generated from the T1-weighted brain volume using infant FreeSurfer’s automatic segmentation code (infant-recon-all; https://surfer.nmr.mgh.harvard.edu/fswiki/infantFS (Zöllei et al. 2020), (2) a second segmentation was done using both T1- and T2-weighted anatomical images and the brain extraction toolbox (Brain Extraction and Analysis Toolbox, iBEAT, v-2.0 cloud processing, https://ibeat.wildapricot.org/ (Wang et al. 2023), (3) the iBEAT segmentation which was more accurate than the FreeSurfer segmentation was manually corrected to fix segmentation errors (like mislabeled white matter voxels or holes) using ITK-SNAP (http://www.itksnap.org/), and (4) the iBEAT corrected segmentation was reinstalled into FreeSurfer and the resulting segmentation was used for further analyses. An inflated mesh of each infant’s cortical surface was generated from the gray-white matter boundary and used for visualization purposes.

**Figure 1:**
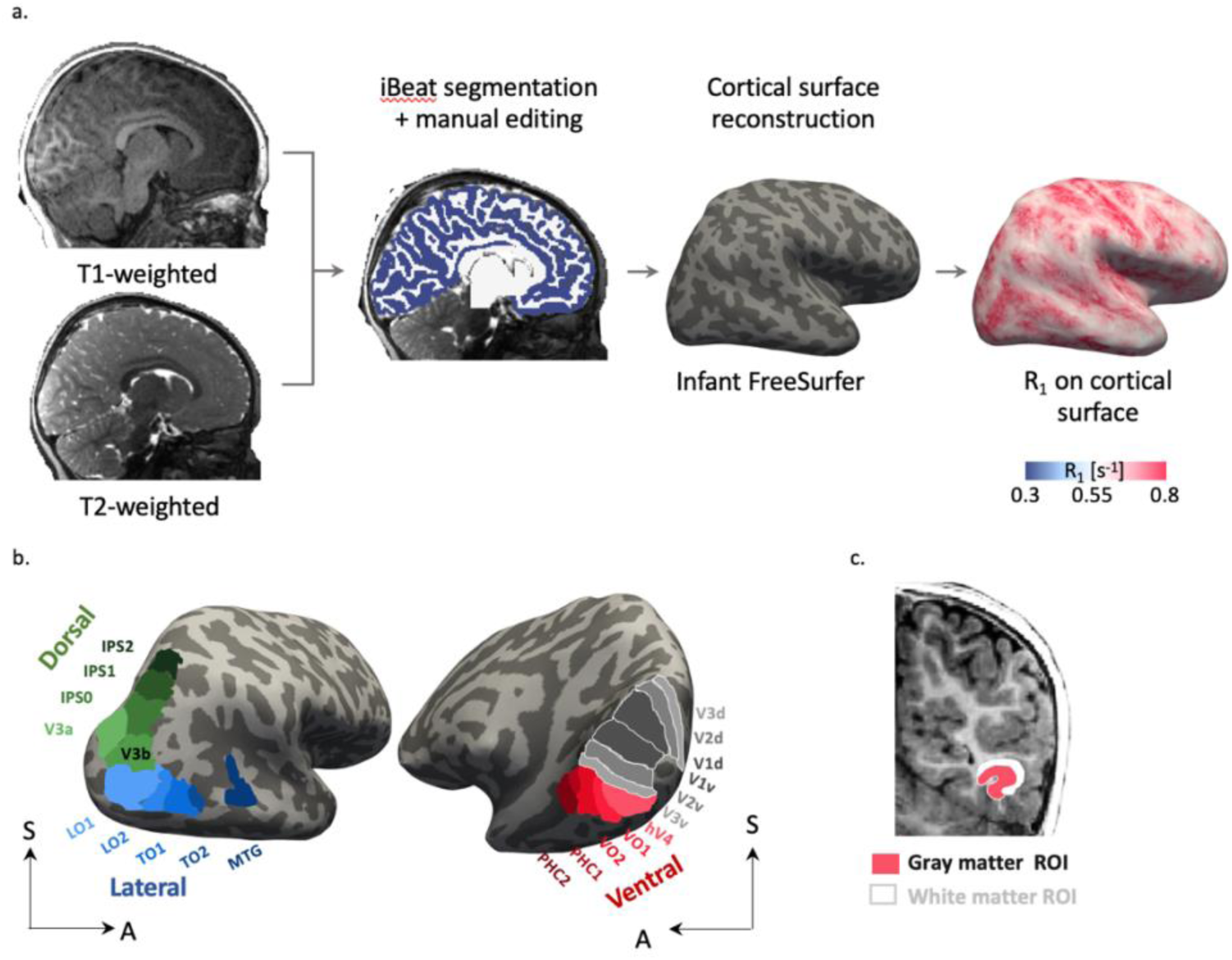
Analysis Pipeline. a) Generation of cortical surfaces using T1- and T2-weighted scans, R_1_ maps, and the projection of R_1_ map on brain surface. b) Regions of interest (ROIs) across the three visual streams on inflated surface maps: dorsal stream (green), lateral stream (blue), and ventral stream (red). We used cortical-based alignment to project the Wang atlas(Wang et al. 2015) to each infant’s cortical surface to delineate ROIs in each infant’s brain. These included the three early visual areas (V1 = V1d + V1v, V2 = V2d + V2v, V3 = V3d + V3v); five regions in the dorsal visual stream (V3a, V3b, IPS0, IPS1, and IPS2), five regions in the lateral visual stream (LO1, LO2, TO1, TO2, and middle temporal gyrus (MTG), delineated using the Rosenke atlas(Rosenke et al. 2021)) and five regions in the ventral visual stream (hV4, VO1, VO2, PCH1, and PCH2). We combined the early visual areas across all three visual streams. c) Coronal slice showing a representative gray matter ROI (pink) and its adjacent white matter (white) used to quantify tissue properties.

#### Delineation of early, dorsal, lateral, and ventral visual areas

Our goal was to examine developmental changes in quantitative R_1_ in the gray and white matter of early visual cortex as well as across multiple regions of interest (ROI) in the dorsal, lateral, and ventral visual processing streams from birth to 1 year of age. To delineate these regions in infants, we utilized brain atlases based on the FreeSurfer adult average brain, which were projected onto each infant’s cortical surface at each timepoint using the cortex-based alignment algorithm in infant FreeSurfer. In the absence of functional imaging data, cortex-based alignment provides the most reliable method for localizing brain areas from atlases to individual brains(Fischl et al. 1999). We used the Wang atlas(Wang et al. 2015) to delineate 17 regions of interest (ROIs) in each participant’s brain: three early visual areas (V1 = V1d + V1v, V2 = V2d + V2v, V3 = V3d + V3v), five retinotopic areas in the dorsal visual stream (V3a, V3b, IPS0, IPS1, IPS2), five retinotopic areas in the ventral stream (hV4, VO1, VO2, PHC1, PHC2), and five regions in the lateral stream, four retinotopic areas (LO1, LO2, TO1, TO2) and one higher level body-selective region in the middle temporal gyrus (MTG), which was delineated using the Rosenke atlas(Rosenke et al. 2021)). As V1, V2, and V3 project to all streams, the same early visual data are shown in all the streams; hence, each stream has eight ROIs (**Fig. 1b**).

#### Analysis of mean R_1_ in gray matter and adjacent white matter during development

To examine the development of microstructural properties in the visual streams, we used R_1_ maps per infant and timepoint and calculated mean R_1_ in both the gray matter of the individual ROIs as well as the underlying white matter of each ROI. Mean R_1_ in gray matter was calculated as the average R_1_ of all voxels from the gray-white matter boundary to the pial surface. To calculate the mean R_1_ in white matter, we first projected the gray matter ROI into its neighboring or adjacent white matter (**Fig. 1c**). This was achieved by excluding the gray-white matter boundary and projecting 50% into the deep white matter underlying each ROI using the FreeSurfer function “mri_label2vol” and the “-proj frac” flag. Mean white matter R_1_ was calculated as the average R_1_ of all voxels in this adjacent white matter ROI. All microstructural measures were generated in infant’s native brain space and each infant contributed to a single value per white and gray matter tissue type ROI.

### Quantification and Statistical Evaluation of R_1_ Development

#### R_1_ development with age and ROI across each visual stream

To quantify R_1_ development in the first year of life, we conducted multiple linear mixed models (LMM)(Bolker et al. 2009) using the ‘fitlme’ function in MATLAB version R2020b (MathWorks, Inc.). LMMs were used to model both longitudinal and cross-sectional aspects of our large-scale data. Per LMM, we related mean R_1_ to age and were applied separately for each stream (ventral/dorsal/lateral), hemisphere (left/right), and tissue type (gray/white). To test if R_1_ significantly develops and varies by ROI within a stream: we used the following formula:

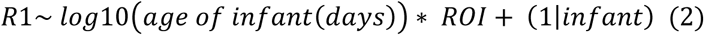

Where age is a continuous variable, ROI is a categorical variable with 8 ROIs per stream, and infant is a repeated measure as we have multiple measurements across the same infant across ROIs and age. In this model, different infants can have different intercepts, but the coefficients of the main factors and their interaction are the same across all infants. As we found significant main effects of age and ROI as well as an interaction between age and ROI (**Supplementary Table 2**), we ran a second LMM separately for each ROI to determine each ROI’s estimated R_1_ at birth (intercept of the LMM) and R_1_ development rate from birth to one year (slope of the LMM):

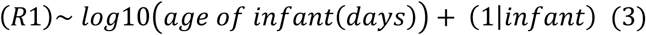

#### R_1_ development across gray and white matter

To test if R_1_ development varies across gray and white matter, we used a third LMM:

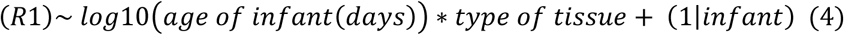

This LMM was run separately for each stream and hemisphere.

#### R_1_ development across streams

To test whether there are developmental differences in R_1_ across the three visual processing streams, we ran the following LMM for both tissue types and hemispheres separately excluding the early visual areas (V1, V2, V3) from each stream as they were common across streams.

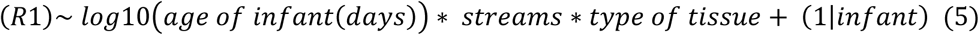

#### Measurement of crossover of white matter values over gray matter

We observed that during the first year of life, R_1_ curves in the white matter cross over those of the gray matter. To quantify the age when this cross-over occurs, we calculated the intersection point of their respective R_1_ curves per ROI per stream. To do this, per tissue type, we first modeled R_1_ as a linear function of log-transformed age to obtain linear fits. The gray and white matter developmental trajectories were expressed as:

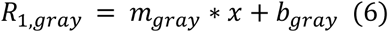

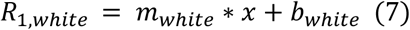

where x represents log_10_ (age in days), and m and b represent the slope and intercept of the fitted line for each tissue type. To find the age at which gray and white matter R_1_ values were equal - i.e., the intersection point – we then solved for x where the two equations intersect:

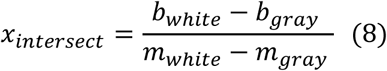

This intersection point in log_10_ (age) was then exponentiated to obtain the corresponding age in days:

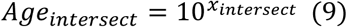

## Results

We first examined R_1_ maps across the brain from birth to one year. Visualizing R_1_ maps of the same infant across four timepoints (newborn, 3 months, 6 months, and 12 months) reveals significant changes in R_1_ from birth to one year (**Fig. 2**). The entire cortex increases in tissue density from approximately 0.4 s^−1^ (in light blue/white) at 17 days to ∼0.7 s^−1^ (in pink/red) at 360 days for this infant indicating large tissue growth (**Fig. 2**). Moreover, R_1_ is not uniformly distributed across cortex at birth. Its development appears to be heterogenous as some areas are redder (higher R_1_) than others at 360 days.

**Figure 2.**
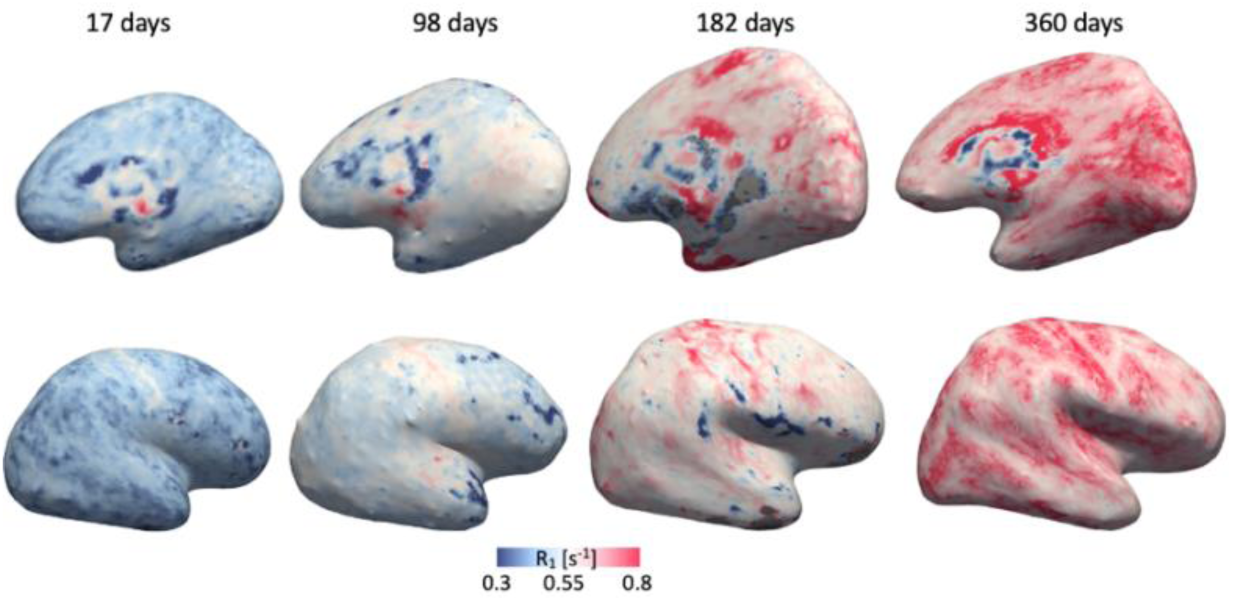
Quantitative R_1_ maps from birth to one year of age. Microstructure tissue density maps R_1_ [s^−1^] in an example infant’s right hemisphere cortical surface scanned across four timepoints (newborn: 17 days; 3 months: 98 days; 6 months: 182 days; one year: 360 days) on medial (top) and lateral views (bottom). Cooler colors (blue) indicate lower R_1_ values, reflecting less dense tissue. There is a progressive increase in R_1_ values from birth to one year (blue, white to pink/red).

Next, we quantified the development of R_1_ across visual areas spanning the three processing streams and tested whether development is uniform or varies as a function of regions and/or streams. We first examine the dorsal visual stream which anatomically lies on the superior portion of the brain, then the lateral stream, and finally the ventral stream, which lies on the inferior portion of the brain.

### How does gray matter tissue microstructure develop across the dorsal visual stream?

To quantity the development of cortical microstructure of dorsal visual stream in the first year of human life, we calculated the mean R_1_ in the gray matter of each of the early visual areas (V1, V2, V3) and dorsal retinotopic areas (V3a, V3b, IPS0, IPS1, IPS2) in each infant and session and plotted mean R_1_ as a function of infants’ age (**Fig. 3a**). We observed that in all ROIs, R_1_ increases in the first year of life with a larger increase in R_1_ from newborn (0-30 days) to six months (∼180 days) than from 6 months to 12 months (**Fig. 3**: right hemisphere (RH); **Supplementary Fig. 2a**: left hemisphere (LH)).

**Figure 3.**
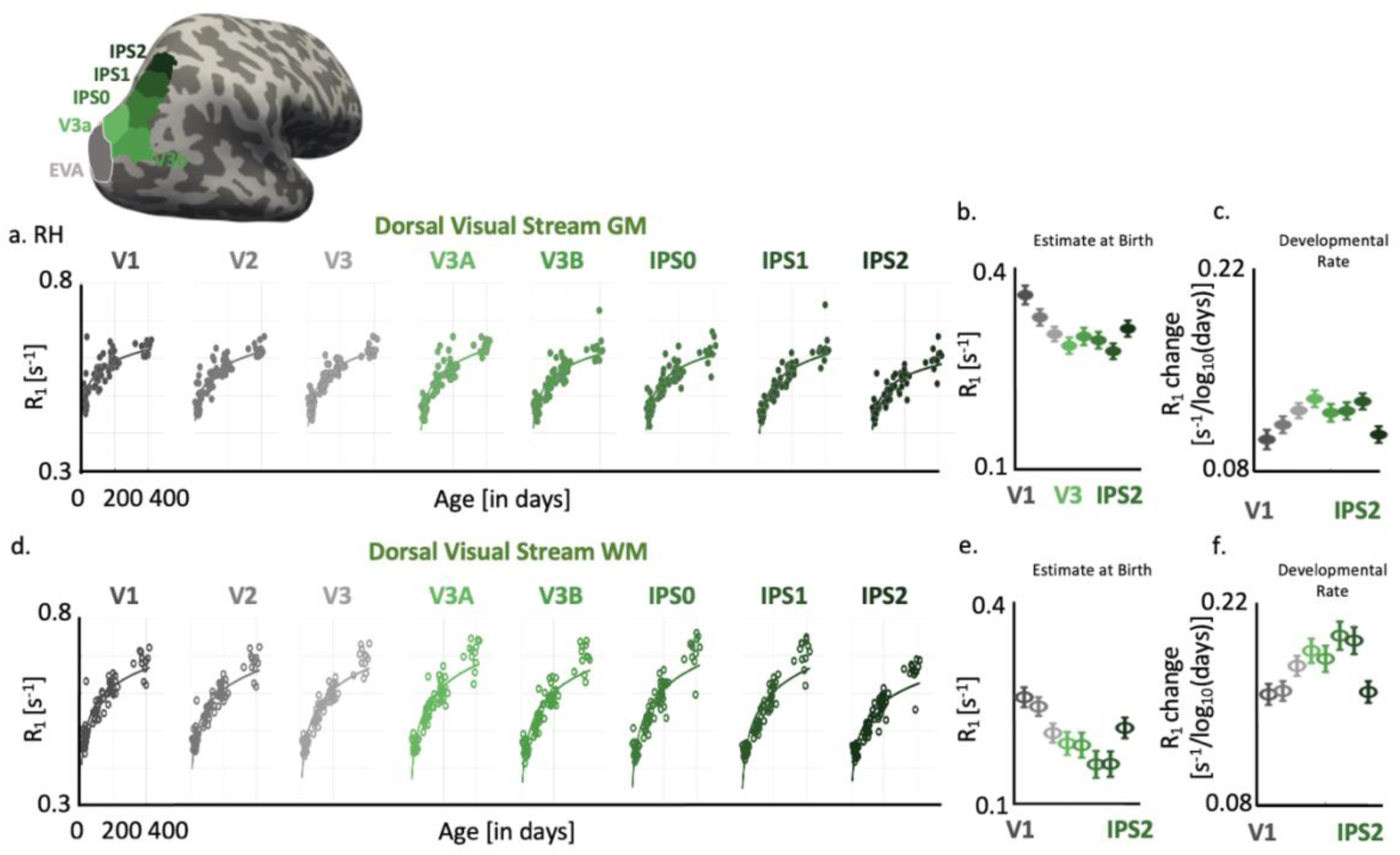
Extensive microstructural tissue growth in the dorsal visual stream from birth to one year of life. a) Inflated cortical surface of an example infant showing the five dorsal regions (in shades of green) combined with early visual areas (in gray). Scatter plots (left to right) show cortical R_1_ as a function of infant’s age per visual area of dorsal stream combined with 3 early visual areas (EVA: V1, V2, V3 + dorsal: V3a, V3b, IPS0, IPS1, and IPS2). Each dot denotes R_1_ per ROI/infant. R_1_ increases longitudinally with faster development in the first 6 months of life than in the later months. Solid lines: Linear mixed model (LMM: (*R*1)∼ *log*10(*age of infant*(*days*)) + (1|*infant*)) estimates of R_1_ development per visual area. b-c) LMM estimates of mean R_1_ at birth (LMM intercept) and R_1_ development (LMM slope). Error bars: standard error on estimates of intercepts and slopes. d-f) Same as in a-c in adjacent white matter ROIs. Data shown are from the right hemisphere. Left hemisphere data are in **Supplementary Fig. 2**

To statistically quantify R_1_ development in the dorsal stream and model both cross-sectional and longitudinal data, we used LMMs (Eq. 2). To account for the nonlinear increase in R_1_, we relate R_1_ to log_10_ (age[days]) and test if age-dependent changes in R_1_ vary across ROIs. We find that: (i) Gray matter R_1_ significantly increases with age in all dorsal visual stream ROIs (main effect of age (RH): *β* = 3.81×10^−4^, SE = 1.90×10^−5^, t_676_ = 20.03, *p* = 2.02×10^−70^, 95% CI = [3.44×10^−4^ 4.19×10^−4^], full statistics for both hemispheres, **Supplementary Table 2**), (ii) Gray matter R_1_ significantly differs across ROIs main effect of ROI (RH): *β* =-6.57×10^−3^, SE = 6.43×10^−4^, t_676_= −10.22, *p* = 6.65×10^−23^, 95% CI = [−7.83×10^−3^ −5.31×10^−3^]), and (iii) Gray matter R_1_ develops differentially across dorsal stream ROIs (significant age by ROI interaction (RH): *β*= 8.68×10^−6^, SE = 3.49×10^−6^, t_676_ = 2.49, *p* = 0.0130, 95% CI = 1.83×10^−6^ 1.55×10^−5^]). R_1_ increases suggest that the dorsal stream gray matter becomes denser with tissue from birth to one year and that different ROIs have different mean tissue density and different developmental trajectories.

Because we found a significant interaction between age and ROI, we used a second LMM relating R_1_ to age to estimate for each ROI R_1_ at birth (LMM intercept) and rate of R_1_ development (LMM slope). We find that R_1_ at birth was higher in early visual areas (RH: **Fig. 3b**, LH: **Supplementary Fig. 2b**). E.g., right V1 has higher R_1_ (mean R_1_±SE: 0.36 ± 0.013[s^−1^]) than later areas like IPS1 (0.27 ± 0.011[s^−1^]). However, the rate of development showed an inverse pattern with higher areas having a faster development rate than earlier areas (RH: **Fig. 3c**, LH: **Supplementary Fig. 2c**). E.g., the rate of R_1_ increase in V1 (0.101 ± 0.006 [s^−1^/log_10_ (age)]) is slower than in IPS1 (0.128 ± 0.0056 [s^−1^/log10 (age)) (RH: **Fig. 3c**, LH: **Supplementary Fig. 2c**, full statistics per ROI in **Supplementary Table 3**). This data shows that across the dorsal visual stream, there is microstructural tissue growth during the first year of life that follows a hierarchical trajectory with early visual areas having higher R_1_ at birth but developing more slowly postnatally than higher order areas up the dorsal visual hierarchy.

### Is there differential development between the gray and white matter in the dorsal visual stream?

We next examined R_1_ development in the superficial white matter adjacent to the dorsal stream visual areas. Like the gray matter, R_1_ in the superficial white matter adjacent to ROIs of the dorsal stream significantly increases with age in a logarithmic manner indicating extensive tissue growth (RH: **Fig. 3d**, LH: **Supplementary Fig. 2d**, main effect of age: RH: *β* = 5.89×10^−4^, SE = 1.92×10^−5^, t_676_ = 30.74, p = 1.63×10^−130^, 95% CI = [5.52×10^−4^ 6.27×10^−4^], full statistics in **Supplementary Table 2**). Mean white matter R_1_ also significantly varied across ROIs (main effect of ROI (RH): *β* = −0.00753, SE = 6.43×10^−4^, t_676_ = −11.71, p = 5.94×10^−29^, 95% CI = [−0.00880 −0.00627]), and R_1_ development significantly varied across ROIs (significant age by ROI interaction (RH): *β* = 1.79×10^−5^, SE = 3.49×10^−6^, t_676_ = 5.12, p = 4.03×10^−7^, 95% CI = [1.10×10^−5^ 2.47×10^−5^]). Additional LMMs per ROI estimated for each region’s superficial white matter R_1_ at birth and development rate (RH: **Figs. 3e-f**, LH: **Supplementary Fig. 2e-f**, full statistics per ROI in **Supplementary Table 4**). As in gray matter, R_1_ in the superficial white matter of early visual areas (e.g., V1) was higher at birth than higher-level regions (e.g., IPS1), but the rate of R_1_ development in the white matter was reversed as white matter R_1_ showed significantly faster increases in higher-level than early visual regions.

We noted that in dorsal stream ROIs the development of R_1_ in the white matter was steeper than in the gray matter (**Figs. 3d-f vs. Figs. 3a-c**) and the estimated R_1_ at birth was lower in the white than gray matter, (RH: **Fig. 3f vs. Fig. 3b**), LH: **Supplementary Fig. 2**). To quantify the differential development across tissue types, per hemisphere, we ran an LMM relating R_1_ to age and tissue type, finding a significant interaction between age and tissue type (age by tissue type interaction (RH): *β* = 0.00024, SE = 1.28×10^−5^, t_1356_ = 18.38, p = 1.35×10^−67^, 95% CI = [0.00021 0.00026], see full statistics in **Supplementary Table 5**). As seen in **Figs. 3a-c**, in newborns, R_1_ in the white matter is lower than R_1_ in the gray matter, but in 12 months, R_1_ in white matter is higher than that in gray matter.

Together data show that gray and white matter R_1_ increases in the first year of life following a hierarchical trajectory across the dorsal visual processing stream, with R_1_ higher at birth in the gray than white matter, but R_1_ developing faster in the white than gray matter.

### Does the lateral stream show a similar pattern of microstructural development as the dorsal stream?

Next, we asked if we see a similar pattern of R_1_ development in the lateral stream as in the dorsal stream. Plotting mean R_1_ as a function of participants’ age in days across early visual areas (V1-V3) and lateral stream ROIs (LO1, LO2, TO1, TO2, MTG) revealed that R_1_ in the gray matter increases in the first year of life following a logarithmic pattern (**Fig. 4a**). Like the dorsal stream, there is larger increase in R_1_ over the first six months of life than the second six (RH: **Fig. 4a**; LH: **Supplementary Fig. 2g**). Quantifying this development, we find that R_1_ in gray matter of the lateral stream significantly increases from birth to one year of age (main effect of age (RH): *β* = 3.80×10^−4^, SE = 2.07×10^−5^, t_676_ = 18.32, *p* = 3.45×10^−61^, 95% CI = [3.39×10^−4^ 4.21×10^−4^], LH and full statistics in **Supplementary Table 2**), and significantly varies across ROIs (main effect of ROI (RH) *β* = −0.00545, SE = 7.01×10^−4^, t_676_= −7.77, *p* = 2.84×10^−14^, 95% CI = [−0.00682 - 0.00407]. R_1_ development significantly varied across ROIs only in the left hemisphere (age by ROI interaction (LH): put here stat of (RH): *β*= 6.49×10^−6^, SE = 3.80×10^−6^, t_676_ = 1.71, *p* = 0.0883, 95% CI = −9.75×10^−7^ 1.40×10^−5^]. Estimating gray matter R_1_ at birth and R_1_ development per ROI, reveals a similar overall pattern like the dorsal stream as early visual areas like V1 have higher R_1_ at birth than higher-order lateral regions like LO. However, unlike the dorsal stream, the hierarchical developmental pattern is less clear (RH: **Figs. 4b-c**, LH: **Supplementary Figs. 2g-i**; full statistics per ROI in **Supplementary Table 6**). In fact, gray matter R_1_ in TO1 and MTG, develops at similar rates to V1, and TO1 also shows similar estimated R_1_ at birth as V2. This finding is interesting as TO1 overlaps with motion-selective hMT(Amano et al. 2009), and prior work suggests that MT in non-human primates is early developing (Bourne and Rosa 2006). Thus, gray matter becomes microstructurally denser in the lateral visual stream during the first year of life, with a heterogeneous but not clearly hierarchical pattern of development.

**Figure 4.**
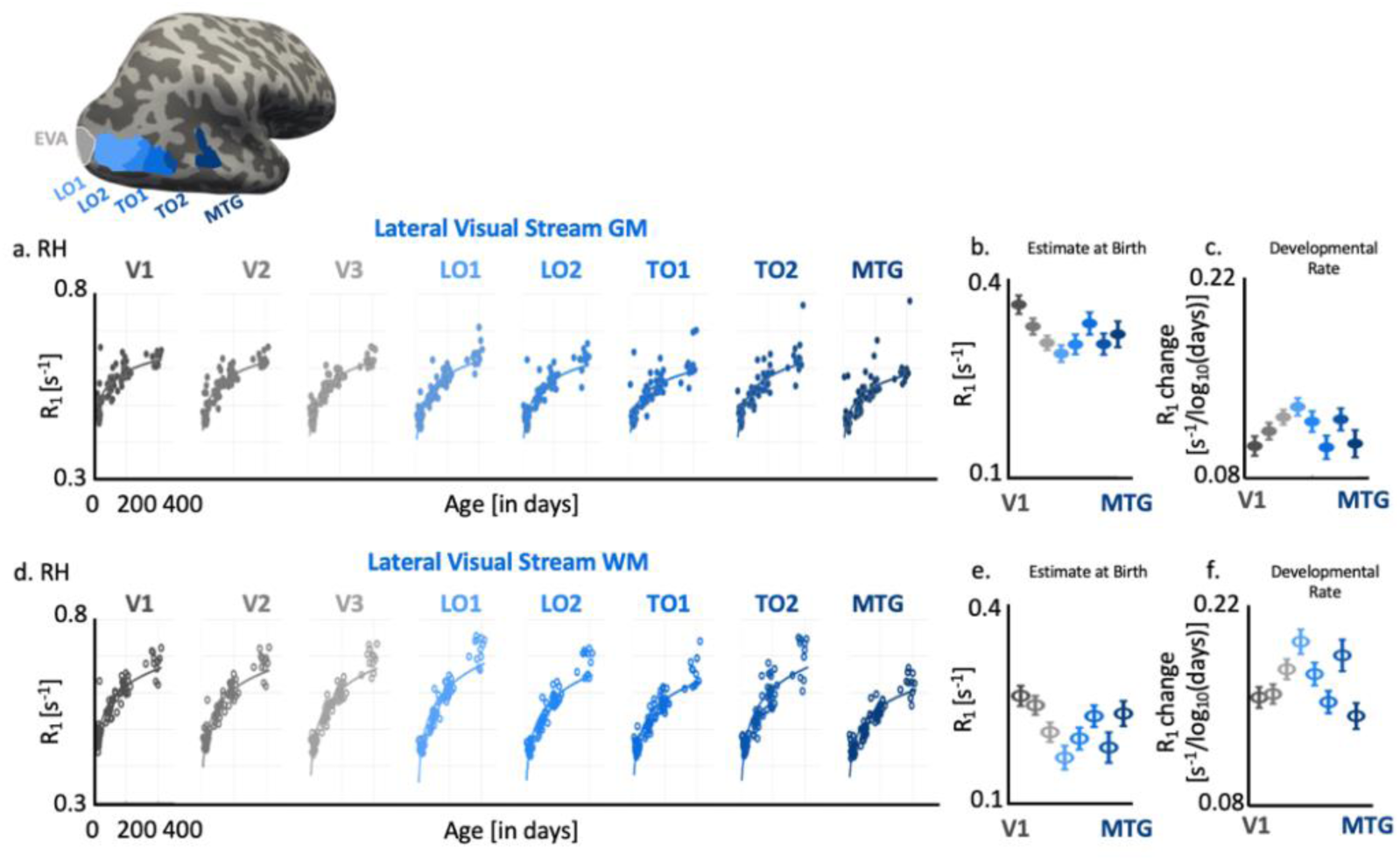
Microstructural tissue growth in the lateral visual stream is hierarchical and heterogeneous from birth to one year of life. a) Inflated cortical surface of an example infant showing the five lateral regions (in shades of blue) combined with early visual areas (in gray). Scatter plots (left to right) showing cortical R_1_ as a function of infant’s age in each visual area of lateral stream combined with three early visual areas (EVA: V1, V2, V3 + lateral: LO1, LO2, TO1, TO2, and MTG). Each dot denotes mean R_1_ per area/infant. R_1_ increases longitudinally with faster development in the first 6 months of life than in the later 6 months. Solid lines: LMM estimates of R_1_ development per visual area. b-c) LMM estimates of mean R_1_ at birth (LMM intercept) and R_1_ development (LMM slope). Error bars: standard error on estimates of intercepts and slopes. d-f) Same as in a-c from adjacent white matter. Data shown are from the right hemisphere. Left hemisphere data are in **Supplementary Fig. 2**.

Likewise, R_1_ in the superficial white matter of the lateral stream significantly increases with age in a logarithmic manner (main effect of age (RH): *β* = 6.28×10^−4^, SE = 2.02×10^−5^, t_676_ = 31.02, *p* = 4.80×10^−132^, 95% CI = [5.88×10^−4^ 6.67×10^−4^], full statistics for both hemispheres in **Supplementary Table 2**) and also varied across ROIs (main effect of ROI (RH): *β* = −0.00646, SE = 6.80×10^−4^, t_676_ = −9.50, p = 3.45×10^−20^, 95% CI = [−0.00780 −0.00513]). Like gray matter, we find differential development of R_1_ in the superficial white matter of the lateral stream only in the left hemisphere (significant age by ROI interaction: (LH): *β* = 1.17×10^−5^, SE = 3.57×10^−6^, t_676_ = 3.26, p = 1.17×10^−3^, 95% CI = [4.63×10^−6^ 1.87×10^−5^]; (RH): *β* = 4.40×10^−6^, SE = 3.69×10^−6^, t_676_ = 1.19, p = 0.233, 95% CI = [−2.84×10^−6^ 1.16×10^−5^], **Figs. 4d-f**, LH: **Supplementary Figs. 2j-l**; full statistics per ROI in **Supplementary Table 7**). In the lateral stream, R_1_ in the white matter is lower at birth than in the gray matter (**Fig. 4e vs. Fig. 4b**) but over the first year of life, R_1_ increases more in the white matter than in the gray matter (**Figs. 4d-f vs. Figs. 4a-c**). Indeed, there is a significantly different development of R_1_ in the gray and white matter of lateral stream ROIs (age by tissue type interaction (RH): *β* = 0.00021 SE = 1.36×10^−5^, t_1356_ = 15.54, p = 3.22×10^−50^, 95% CI = [0.00019 0.00024], full statistics in **Supplementary Table 5**). Like the dorsal stream, the development of white matter R_1_ across ROIs mirrors the heterogeneity of gray matter development but overall develops faster with R_1_ in the white matter exceeding R_1_ in the gray matter in 12-month-olds.

### What is the pattern of microstructural tissue development in the human ventral visual stream?

Given that we see a clearer hierarchical pattern of R_1_ development in the dorsal than lateral visual stream, we next asked if the development of the ventral visual cortex is similar to one of these streams or follows its own trajectory. Results revealed a development pattern similar to that of the dorsal stream. Gray matter R_1_ significantly increases in the first year of life in a logarithmic pattern (RH: **Fig. 5a**: LH: **Supplementary Fig. 2m**, main effect of age (RH): *β* = 3.97×10^−4^, SE = 1.89×10^−5^, t_676_ = 21.01, *p* = 7.90×10^−76^, 95% CI = [3.60×10^−4^ 4.34×10^−4^] full statistics in **Supplementary Table 2)**. Gray matter R_1_ also varies significantly across ROIs (RH: β = −0.00447, SE = 6.34×10^−4^, t_676_ = −7.06, p = 4.09×10^−12^, 95% CI = [−0.00572 - 0.00323]) and develops differentially across ROI (significant age by ROI interaction (RH) β = 9.92×10^−6^, SE = 3.44×10^−6^, t_676_ = 2.89, p = 0.00399, 95% CI = [3.18×10^−6^ 1.67×10^−5^]). Additional LMMs (Eq. 3) estimating gray matter R_1_ at birth and developmental rate across ROIs revealed that early visual areas like V1 are denser at birth than higher-level areas for instance PHC1/PHC2. However, the rate of R_1_ development showed a reversal as higher-level regions like PHC1/PHC2 showed steeper increases in R_1_ than V1 (RH: **Figs. 5b-c**, LH: **Supplementary Figs. 2n-o**; full statistics per ROI in **Supplementary Table 8**). This reversal between initial maturity and growth rate supports a hierarchical and heterogeneous pattern of development similar to our observations in the dorsal stream.

**Figure 5.**
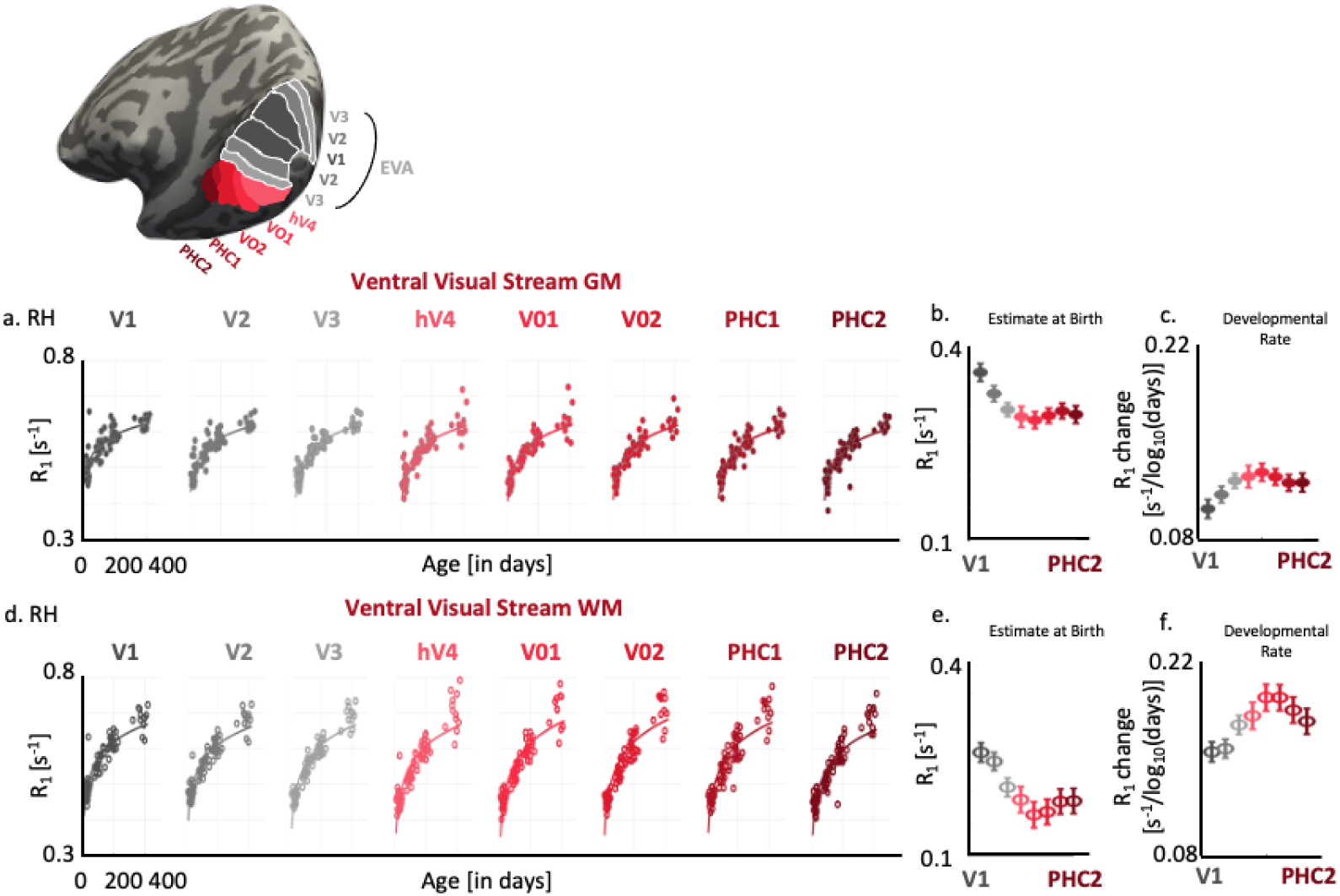
Ventral stream undergoes hierarchical and heterogeneous tissue growth from birth to one year of life. a) Inflated cortical surface of an example infant showing the five ventral regions (in shades of red) combined with early visual areas (in gray). Scatter plots (left to right) showing cortical R_1_ as a function of infant’s age in each visual area of ventral stream combined with three early visual areas (EVA: V1, V2, V3 + ventral: hV4, VO1, VO2, PHC1, and PHC2). Each dot denotes mean R_1_ per area/infant. R_1_ increases longitudinally with faster development in the first 6 months of life than in the later 6 months. Solid lines: Linear mixed model estimates of R_1_ development per visual area. b-c) LMM estimates of mean R_1_ at birth (LMM intercept) and R_1_ development (LMM slope). Error bars: standard error on estimates of intercepts and slopes. d-f) Same as in a-c from adjacent white matter. Data shown are from the right hemisphere. Left hemisphere data are in **Supplementary Fig. 2**.

Examining the development of the adjacent white matter in the ventral stream also revealed that R_1_ increases with age (main effect of age: RH: β = 6.22×10^−4^, SE = 1.79×10^−5^, t_676_ = 34.72, p = 2.23×10^−152^, 95% CI = [5.87×10^−4^ 6.57×10^−4^]), that R_1_ varies across ROIs (β = −0.0058, SE = 5.93×10^−4^, t_676_ = −9.78, p = 3.18×10^−21^, 95% CI =[−0.00696 −0.00464]) and that R_1_ development significantly varies across ROIs (significant interaction between age and ROI, β = 2.12×10^−5^, SE = 3.22×10^−6^, t_676_ = 6.59, p = 8.63×10^−11^, 95% CI = [1.49×10^−5^ 2.75×10^−5^]; RH: **Fig. 4d**, LH: **Supplementary Fig. 2p**; full statistics in **Supplementary Table 2**). In the white matter as well, there is a clear hierarchical pattern of development (RH: **Fig. 4e-f**, LH: **Supplementary Figs. 2q-r**; full statistics per ROI in **Supplementary Table 9**). These observations reveal that microstructural tissue development in the ventral visual stream is both heterogeneous and hierarchical.

Finally, there were significant developmental differences between the ventral gray and white matter as white matter develops faster than the gray matter (significant age and type of tissue interaction (RH: *β* = 2.5 ×10^−4^, SE = 1.16×10^−5^, t_1356_ = 21.136, p = 6.10×10^−86^, 95% CI = [2.2 ×10^−4^ 2.6 ×10^−4^]) in addition to a significant main effects of age and tissue type (|ts|> 10.23, ps < 1.03×10^−23^; for full statistics see **Supplementary Table 5**). These results indicate that gray and white matter along the ventral stream follow different developmental trajectories, with white matter developing faster than cortex.

### Are there differential developments between the three visual streams?

Given that we observed a clearer hierarchical developmental pattern in visual areas of the dorsal and ventral stream than that of the lateral stream, we asked if microstructural development during the first year of life varies across the three streams. In this analysis, we excluded early visual areas (V1, V2, V3) which are shared across streams. We did not find significant difference in R_1_ across streams (no main effect of stream: right: *β* = 0.0048, SE = 0.0033, t_2542_ = 1.4633, p = 0.14, 95% CI = −0.0016 0.011, left: *β* = 0.0036, SE =0.0031, t_2542_ = 1.15, p = 0.24, 95% CI = [−0.0025 0.0097] no significant differential development across streams (no significant age by stream or age by stream and tissue interactions (−0.77>ts>0.07, *ps*>0.44), LMM: ((*R*1)∼ *log*10(*age of infant*(*days*)) ∗ *streams* ∗ *type of tissue* + (1|*infant*)).

Finally, as we observed that in all streams, white matter R_1_ in newborns is lower than gray matter R_1_ but white matter R_1_ in 1-year-olds is higher than gray matter R_1_, we sought to estimate the age at which R_1_ in the white matter catches up with R_1_ in the gray matter. We refer to this point as the crossover age. We found that this crossover is around 2 months of age, and occurs earliest in the dorsal stream (RH: Mean_crossoveage_ ± SD_crossoverage_: 57.01 ± 6.65 days; LH: 57.27 ± 7.27 days), then in the ventral stream (RH: Mean_crossoveage_ ± SD_crossoverage_: 58.69 ± 7.25 days; LH: 62.39 ± 15.08 days), and latest in the lateral stream (RH: Mean_crossoveage_ ± SD_crossoverage_: 68.41 ± 17.46 days; LH: 60.92 ± 7.91 days) (**Supplementary Fig. 3**). Together results suggest that the rate and sequence of microstructural development in the gray and adjacent white matter are relatively similar across the dorsal, ventral, and lateral visual streams in the first year of life.

## Discussion

Using cross-sectional and longitudinal infant data, we provide the first systematic analysis of microstructural changes in gray matter and the adjacent white matter during the first year of human life. Our findings reveal extensive microstructural tissue growth across all three visual processing streams during the first year of life, with similar developments across streams. Tissue growth is faster in the first six months and is followed by a slower rate of development in the second half of the year. Furthermore, the development of microstructure in both gray and adjacent white matter of dorsal and ventral visual streams is hierarchical and heterogeneous, as primary visual areas, such as V1, have denser tissue at birth but develop more slowly than higher-level visual areas postnatally. White matter develops faster than gray matter postnatally, but gray matter is surprisingly denser than white matter at birth. Overall, our findings shed light on how tissue properties along visual hierarchies mature after birth and provide the necessary testbed for studying gray and white matter maturation in other sensory and cognitive domains.

### Logarithmic trajectory of microstructural tissue properties in the first year of human life

R_1_ development in the visual system follows a non-linear, logarithmic pattern in the first year. Our prior work showed that R_1_ develops linearly in the first six months of life, however, R_1_ development begins to decelerate in the later months. This logarithmic pattern may reflect a period of rapid structural development that may be required for emerging sensory processes early in life(Rakic 2009). However, by the second half of the first year, as infants develop specialized behaviors - like better motion tracking, increased visual attention, or emergence of visual category representations, microstructural maturation might shift toward refining, stabilizing, or even pruning of circuits as is seen during childhood and adolescence. The slowing of tissue growth in the second half of the year might also reflect the brain becoming more selective(Kosakowski et al. 2022, 2023) as it reinforces existing pathways to support sensory experiences and salient stimuli. Interestingly, this logarithmic pattern of R_1_ development is also consistent with other macrostructural changes during infancy, such as increases in gray and white matter volume, cortical thickness, and surface area(Bethlehem et al. 2022; Ahmad et al. 2023; Tung et al. 2025), suggesting that rates of macro- and micro-level changes may be linked to support emerging behaviors. Detecting these subtle but impactful shifts in development is only possible with longer developmental windows, and our results reveal the importance of looking beyond early infancy to study how anatomy and microstructure unfold over time.

### Hierarchical and heterogeneous tissue growth in gray and white matter of visual streams

Extending our observation window across one year and multiple areas of the three visual hierarchies allowed us to examine the extent of tissue growth at birth and the sequence and rate of development within the visual system. In dorsal and ventral visual streams, there is clear hierarchical and heterogeneous development: early visual areas like V1 are more mature or denser with microstructural tissue at birth but develop slower than higher-level regions ascending the visual hierarchies. We find that this pattern was found not only in cortex but also in the adjacent white matter. This result is novel because it shows that cortex and underlying white matter are not uniform at birth. Even within the three areas of early visual cortex, V1 - V2 - V3, there is a clear high-to-low change in R_1_ values at birth, with further lowering of R_1_ in the higher-level areas (ventral: V01/VO2/PHC1; dorsal: IPS0-IPS1). In contrast, R_1_ development is progressively faster ascending visual hierarchies. In the lateral stream, although development is heterogeneous across its areas, the hierarchical pattern is less clear than that in dorsal and ventral visual streams. Theoretically, the trajectory of hierarchical microstructural growth reflects an interplay between genetics and sensory experiences after birth. V1, for example, lies in the calcarine sulcus which is an early emerging sulcus in utero (between the 16^th^ to 19^th^ gestational week)(Chi et al. 1977; Nishikuni and Ribas 2013), and has some level of retinotopic functional connectivity and organization by infancy in both primates (Arcaro and Livingstone 2017) and humans (Ellis et al. 2021). Whereas higher-level retinotopic areas like PHC1, for example, which lies in the collateral sulcus(Stenger et al. 2022) a sulcus that emerges after the calcarine in utero(Chi et al. 1977), shows prolonged development in infancy and childhood(Golarai et al. 2007; Natu et al. 2021a; Tung et al. 2025). We hypothesize that the more mature V1 at birth may provide a scaffolding for maturation of higher-level areas after birth but the diverse and rich postnatal visual environment may accelerate activity-dependent mechanisms like myelination(Miller et al. 2012), synaptic growth(Katz and Shatz 1996), and dendritic arborization(Elston and Fujita 2014) in higher-level areas. Hence, hierarchical organization might be a fundamental characteristic of the visual system as new processes develop and evolve from older ones(Barrett 2012), raising questions about whether this is a general organizing principle of sensory processes, like the auditory or somatosensory cortex. Overall, our findings provide the first comprehensive measurement of microstructural tissue growth in a sensory cortex during the first year of life and raise the hypothesis that if microstructural maturation is linked to functional specialization, it may follow a similar hierarchical trajectory.

### Lateral visual stream develops in a heterogeneous but not in a hierarchical manner

Even though we do not observe significant developmental differences between the three visual streams, we find subtle variations between the dorsal/ventral versus lateral streams. Specifically, a clear hierarchical developmental pattern from early to high-level areas is less prominent in the lateral stream. In fact, two higher-level areas, TO1 and MTG develop at similar rates to V1, and TOI also has similar R_1_ values as V2 at birth. This result is intriguing as the lateral surface of the temporal–occipital (TO) boundary and areas TO1/TO2(Amano et al. 2009) are known to overlap with the motion-selective cortex hMT+, which is posited to develop in early infancy(Biagi et al. 2023). This result suggests that development is heterogeneous but may not be hierarchical along the lateral stream hierarchy. Furthermore, we also observed early signs of lateralization in the lateral stream: there is a significant interaction between age and ROI in the left but not in the right lateral stream. Given that the lateral stream is associated with social communication and dynamic visual functions that are typically left-lateralized later in life, this asymmetry may reflect early structural foundations of functional lateralization later in childhood.

### White matter develops faster than gray matter, postnatally

Prior studies have examined white matter development during infancy(Deoni et al. 2011; Dubois et al. 2014; Grotheer et al. 2022; Kubota et al. 2025), however, their focus is primarily on long-range, white matter fiber tracts. These tracts may or may not align with local cortical regions or even impact cortical development in individual ROIs. By focusing on localized microstructure in white matter, just beneath each visual ROI, we captured how local white matter and its adjacent cortical tissue develop in tandem. Surprisingly, in all visual streams, gray matter is denser with tissue than white matter at birth, but white matter develops faster than gray matter postnatally. Gray matter mainly contains glia and neuronal processes like cell bodies, and the process of neurogenesis and migration is thought to be completed within the first few weeks of postnatal life(Rakic 2009; Clowry et al. 2010) resulting in a significant increase in cortical thickness and surface area(Bethlehem et al. 2022). One theoretical explanation may be that the cortex gets a head start in utero, and the high tissue density is reflected in the higher values in gray than white matter R_1_ at birth. However, the process of myelination of the white matter is thought to be activity-dependent(Fields 2015) and occurs mainly postnatally in the visual system via sensory experiences. Finding that white matter surpasses gray matter in R_1_ values by around two months of age reflects this shift in tissue densities as white matter becomes denser with myelin to support cortical processing. The differential development of local gray and white matter tissue may not only play a role in functional development but also in cortical folding(Richman et al. 1975; Ronan et al. 2014), which can be explored in future work.

Our findings open promising novel avenues for future research as much remains to be uncovered about the relationship between microstructural growth and functional development. Using fMRI, future studies can explore how changes in microstructure relate to emerging visual(Deen et al. 2017; Ellis and Turk-Browne 2018; Kosakowski et al. 2022) and cognitive functions(Yates et al. 2025) in infancy. For instance, investigating whether regions that show faster gray matter development also undergo faster functional development could help clarify how tissue growth supports functional and behavioral milestones. By combining anatomical and functional data, researchers can build comprehensive and integrative models of human brain development, targeting when critical changes occur, and how developments vary across sensory and cognitive domains. Finally, extending MRI tracking beyond the first year of life will be informative for identifying sensitive periods of development and for understanding how early brain changes scaffold later abilities such as language, attention, and social perception.

In conclusion, this study offers a comprehensive examination of how the infant visual system develops over the first year of life by examining both gray and white matter microstructure across multiple, functionally diverse streams. By extending the developmental window across the entire first year of life, we identified key transitions in growth patterns and shifts that may reflect changes in experience-dependent refinement. Overall, our study has important implications for functional and behavioral development of the infant visual system.

## Acknowledgments

This research was supported by NIH grants R01EY033835, R21 EY030588, Stanford Wu Tsai Neurosciences Institute Big Ideas Grant Phase I, and Stanford Wu Tsai Neurosciences Institute Accelerator grants to KGS.

## Contributions

K.P: participant recruitment, data acquisition, data preprocessing, statistical analysis, manuscript writing; A.A: data preprocessing, statistical analysis, C.T, S.S.T, B.F, X.Y: participant recruitment, data acquisition, data preprocessing; D.O: data acquisition and preprocessing; J.H: data preprocessing; H.W. sequence development; K.G.S and V.S.N. designed, oversaw all components of the study and data analyses, and wrote the manuscript. All co-authors read and approved the submitted manuscript.

## Competing interests

The authors declare no competing interests.

## Data and code availability

Source data and codes used in the analyses and to reproduce figures are available on the GitHub repository: https://github.com/VPNL/visualstreamdevelopment. Requests for raw data should be directed to the corresponding author, Vaidehi S. Natu.

---

**Supplementary Figure 1.**
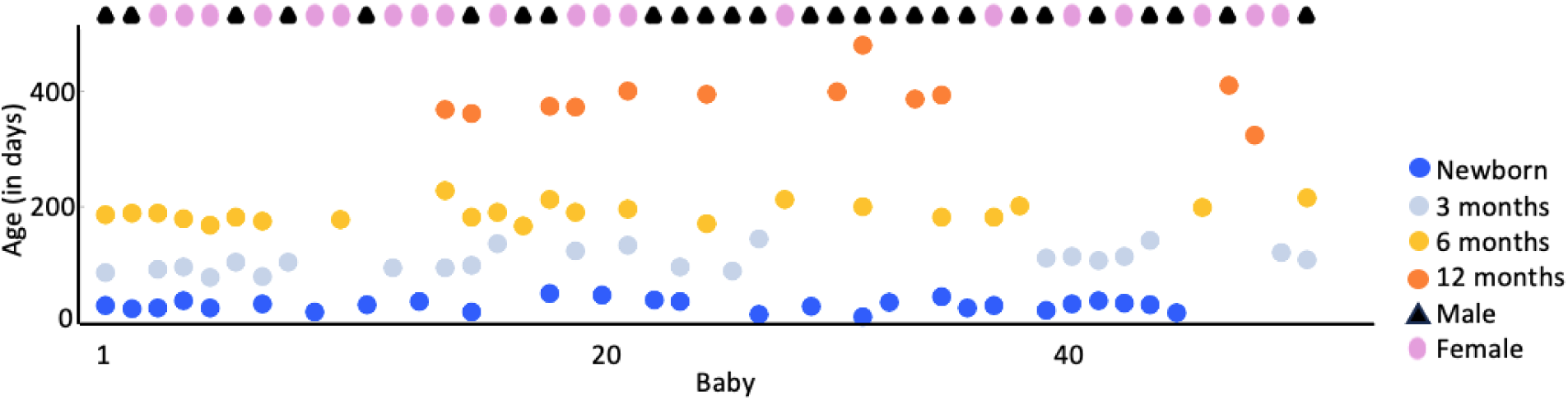
Longitudinal infant data. Scatterplot showing longitudinal data per subject on the x-axis across 4 different four time points (newborn (dark blue), 3 months (light blue), 6 months (light orange), 12 months (dark orange, each triangle represents: male, each oval represents: female). For example, baby 1 has MRI data at newborn, 3 months, and 6 months. Age (in days) is age at the time of their MRI sessions. (See full demographic table in **Supplementary Table 1)**

**Supplementary Figure 2.**
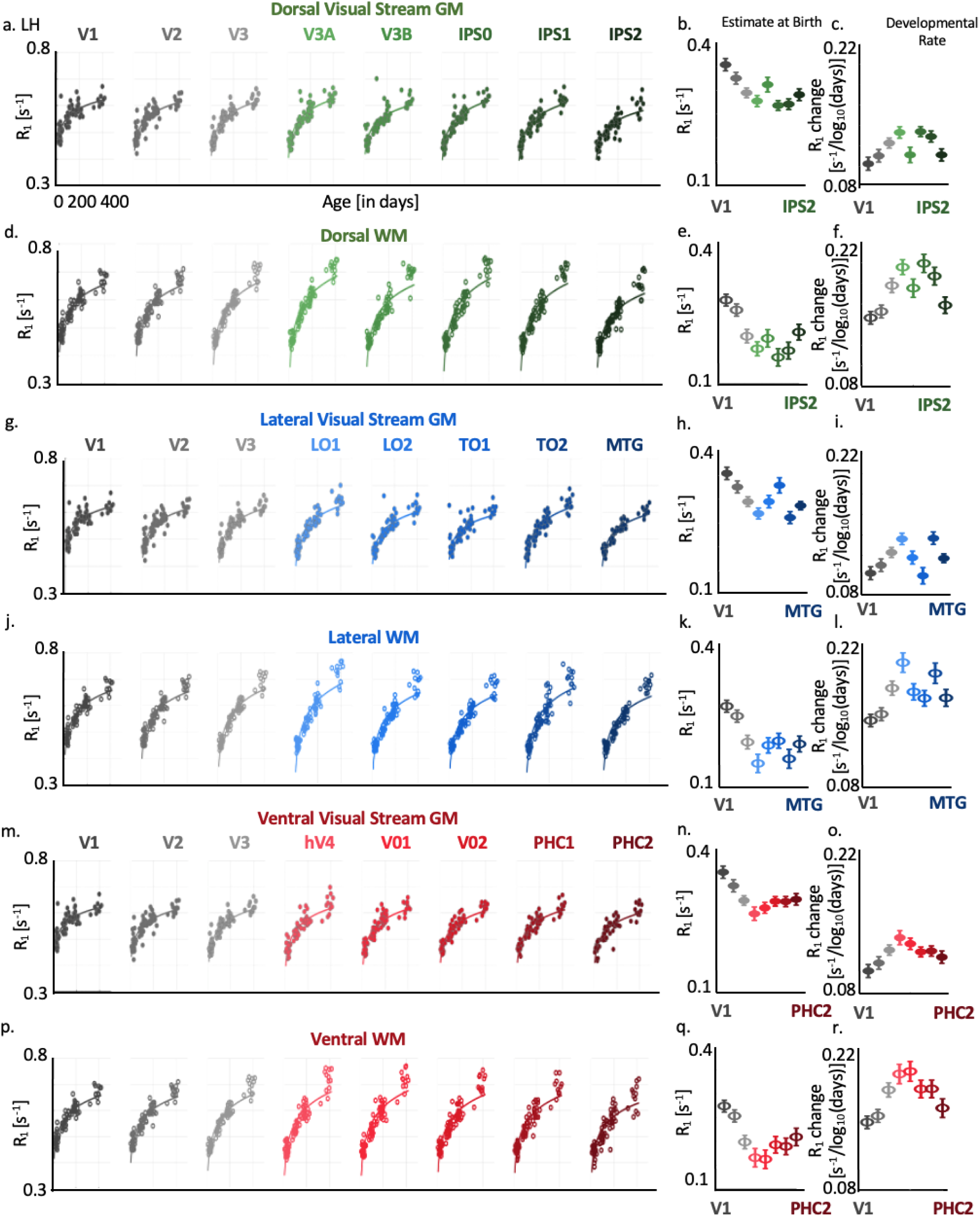
Tissue growth in gray and white matter of left dorsal, lateral, and ventral visual streams. **(a)** R_1_[s^−1^] increases logarithmically in the first year of life. *Dot:* mean R_1_ per ROI, per infant. *Line:* Log fit. (b-c) Estimate of R_1_ at birth and rate of R_1_ development using LMM relating R_1_ versus log_10_ (age in days). *Error bars:* standard error on estimates of slopes and intercept. (d-r) Same as in a-c for other GM and WM visual areas in the left hemisphere.

**Supplementary Figure 3.**
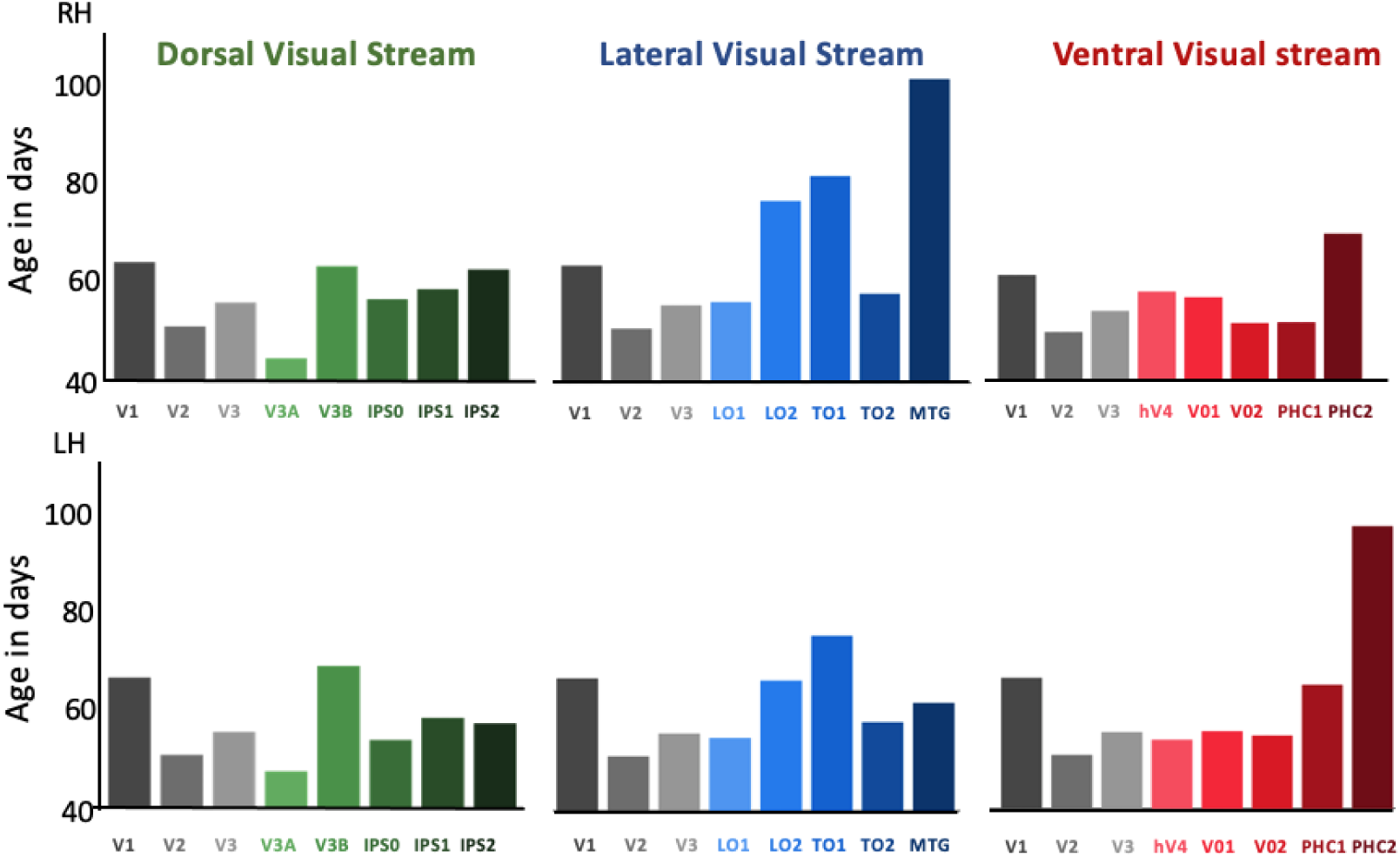
Crossover ages when white matter R_1_ values increase more than gray matter R_1_ in all visual streams. Top row shows right hemisphere and bottom row shows left hemisphere ages in dorsal, lateral, and ventral visual streams respectively. *LH/RH:* Left/right hemisphere.

**Supplementary Table 1.** Participant Demographics. Table provides participant demographics, including their birth sex, ethnicity, and their age in days at the time of their MRI sessions at four time points (newborn, 3 months, 6 months, 12 months). Dash (’-’) indicates that a session did not occur for a participant at that specific time point or participant failed to stay still during anatomical or quantitative scans.

| Subj. No | Birth Sex | Ethnicity | Newborns | 3-month-olds | 6-month-olds | 1-year-olds |
| --- | --- | --- | --- | --- | --- | --- |
| 1 | M | White | 29 | 85 | 185 | - |
| 2 | M | White | 23 | - | 189 | - |
| 3 | F | White | 24 | 91 | 189 | - |
| 4 | F | White | 37 | 95 | 179 | - |
| 5 | F | Asian & White | 24 | 78 | 167 | - |
| 6 | M | Hispanic | - | 104 | 181 | - |
| 7 | F | White | 31 | 79 | 174 | - |
| 8 | M | Asian | - | 104 | - | - |
| 9 | F | Asian | 18 | - | - | - |
| 10 | F | Asian & White | - | - | 177 | - |
| 11 | M | White | 30 | - | - | - |
| 12 | F | Asian & White | - | 94 | - | - |
| 13 | F | White | 35 | - | - | - |
| 14 | F | Asian | - | 94 | 227 | 368 |
| 15 | M | White | 17 | 98 | 182 | 360 |
| 16 | F | Asian & White | - | 135 | 190 | - |
| 17 | M | White | - | - | 166 | - |
| 18 | M | Asian & White | 50 | - | 212 | 373 |
| 19 | F | White | - | 123 | 190 | 372 |
| 20 | F | Asian | 47 | - | - | - |
| 21 | F | Asian & White | - | 133 | 195 | 399 |
| 22 | M | White | 38 | - | - | - |
| 23 | M | Asian & White | 35 | 96 | - | - |
| 24 | M | White | - | - | 170 | 393 |
| 25 | M | Asian & White | - | 88 | - | - |
| 26 | M | White | 14 | 144 | - | - |
| 27 | F | White | - | - | 212 | - |
| 28 | M | Hispanic | 27 | - | - | - |
| 29 | M | White & Black | - | - | - | 398 |
| 30 | M | Asian & White | 9 | - | 199 | 479 |
| 31 | M | White | 34 | - | - | - |
| 32 | M | Asian | - | - | - | 385 |
| 33 | M | White | 44 | - | 182 | 392 |
| 34 | M | Asian & White | 24 | - | - | - |
| 35 | F | Hispanic & White | 29 | - | 181 | - |
| 36 | M | White | - | - | 201 | - |
| 37 | M | Asian | 21 | 111 | - | - |
| 38 | F | Hispanic & White | 31 | 113 | - | - |
| 39 | M | White | 37 | 106 | - | - |
| 40 | F | White | 33 | 113 | - | - |
| 41 | M | White | 30 | 141 | - | - |
| 42 | M | Asian & White | 16 | - | - | - |
| 43 | F | White | - | - | 198 | - |
| 44 | M | White | - | - | - | 409 |
| 45 | F | White | - | - | - | 323 |
| 46 | F | White | - | 121 | - | - |
| 47 | M | Hispanic & White | - | 108 | 214 | - |

**Supplementary Table 2.** Statistical significance per linear mixed model (LMM). LMMs quantifying the relationship between microstructure (R_1_ [s^−1^]) and age [log_10_ (age in days)] across the 8 regions of interest (ROls) per stream (dorsal, lateral, and ventral) and tissue type (gray matter(GM)/white matter (WM)). LMM: *R1 ∼ log10(age of infant) × ROI + (1*|*Infant).* Three effects are shown per model per hemisphere: main effects of age arnd ROls, and interaction between age and ROI. Statistics: *SE:* Standard error; *DF:* degrees of freedom; *tstats:* t-statistics; *CI =* confidence in,tervals. *LH/RH:* left/right hemisphere.

| Stream/<br>Tissue Type | Hemi | Effect | $\beta$ Estimate | SE | DF | tStats | pValue | CI lower | CI upper |
| --- | --- | --- | --- | --- | --- | --- | --- | --- | --- |
| Dorsal GM | LH | Age | 3.65e-04 | 1.97e-05 | 676 | 18.49 | 4.52e-62 | 3.26e-04 | 4.04e-04 |
|  | LH | ROI | -7.14e-03 | 6.66e-04 | 676 | -10.72 | 6.99e-25 | -8.45e-03 | -5.83e-03 |
|  | LH | Age x ROI | 1.25e-05 | 3.61e-06 | 676 | 3.47 | 5.51e-04 | 5.44e-06 | 1.96e-05 |
|  | RH | Age | 3.81e-04 | 1.90e-05 | 676 | 20.03 | 2.02e-70 | 3.44e-04 | 4.19e-04 |
|  | RH | ROI | -6.57e-03 | 6.43e-04 | 676 | -10.22 | 6.65e-23 | -7.83e-03 | -5.31e-03 |
|  | RH | Age x ROI | 8.68e-06 | 3.49e-06 | 676 | 2.49 | 1.30e-02 | 1.83e-06 | 1.55e-05 |
| Dorsal WM | LH | Age | 5.76e-04 | 1.78e-05 | 676 | 32.37 | 1.61e-139 | 5.41e-04 | 6.11e-04 |
|  | LH | ROI | -7.76e-03 | 5.97e-04 | 676 | -12.98 | 1.41e-34 | -8.93e-03 | -6.58e-03 |
|  | LH | Age x ROI | 2.00e-05 | 3.24e-06 | 676 | 6.19 | 1.06e-09 | 1.37e-05 | 2.64e-05 |
|  | RH | Age | 5.89e-04 | 1.92e-05 | 676 | 30.74 | 1.63e-130 | 5.52e-04 | 6.27e-04 |
|  | RH | ROI | -7.53e-03 | 6.43e-04 | 676 | -11.71 | 5.94e-29 | -8.80e-03 | -6.27e-03 |
|  | RH | Age x ROI | 1.79e-05 | 3.49e-06 | 676 | 5.12 | 4.03e-07 | 1.10e-05 | 2.47e-05 |
| Lateral GM | LH | Age | 3.72e-04 | 1.94e-05 | 676 | 19.11 | 1.99e-65 | 3.33e-04 | 4.10e-04 |
|  | LH | ROI | -6.21e-03 | 6.59e-04 | 676 | -9.43 | 6.57e-20 | -7.51e-03 | -4.92e-03 |
|  | LH | Age x ROI | 1.17e-05 | 3.57e-06 | 676 | 3.26 | 1.17e-03 | 4.63e-06 | 1.87e-05 |
|  | RH | Age | 3.80e-04 | 2.07e-05 | 676 | 18.32 | 3.45e-61 | 3.39e-04 | 4.21e-04 |
|  | RH | ROI | -5.45e-03 | 7.01e-04 | 676 | -7.77 | 2.84e-14 | -6.82e-03 | -4.07e-03 |
|  | RH | Age x ROI | 6.49e-06 | 3.80e-06 | 676 | 1.71 | 8.83e-02 | -9.75e-07 | 1.40e-05 |
| Lateral WM | LH | Age | 5.82e-04 | 1.85e-05 | 676 | 31.45 | 1.90e-134 | 5.46e-04 | 6.18e-04 |
|  | LH | ROI | -7.15e-03 | 6.22e-04 | 676 | -11.50 | 4.59e-28 | -8.37e-03 | -5.93e-03 |
|  | LH | Age x ROI | 1.79e-05 | 3.37e-06 | 676 | 5.32 | 1.39e-07 | 1.13e-05 | 2.46e-05 |
|  | RH | Age | 6.28e-04 | 2.02e-05 | 676 | 31.02 | 4.80e-132 | 5.88e-04 | 6.67e-04 |
|  | RH | ROI | -6.46e-03 | 6.80e-04 | 676 | -9.50 | 3.45e-20 | -7.80e-03 | -5.13e-03 |
|  | RH | Age x ROI | 4.40e-06 | 3.69e-06 | 676 | 1.19 | 2.33e-01 | -2.84e-06 | 1.16e-05 |
| Ventral GM | LH | Age | 3.85e-04 | 1.94e-05 | 676 | 19.85 | 1.86e-69 | 3.47e-04 | 4.23e-04 |
|  | LH | ROI | -4.70e-03 | 6.56e-04 | 676 | -7.17 | 1.97e-12 | -5.99e-03 | -3.41e-03 |
|  | LH | Age x ROI | 9.13e-06 | 3.56e-06 | 676 | 2.57 | 1.04e-02 | 2.15e-06 | 1.61e-05 |
|  | RH | Age | 3.97e-04 | 1.89e-05 | 676 | 21.01 | 7.90e-76 | 3.60e-04 | 4.34e-04 |
|  | RH | ROI | -4.47e-03 | 6.34e-04 | 676 | -7.06 | 4.09e-12 | -5.72e-03 | -3.23e-03 |
|  | RH | Age x ROI | 9.92e-06 | 3.44e-06 | 676 | 2.89 | 3.99e-03 | 3.18e-06 | 1.67e-05 |
| Ventral WM | LH | Age | 6.10e-04 | 1.95e-05 | 676 | 31.23 | 3.16e-133 | 5.72e-04 | 6.49e-04 |
|  | LH | ROI | -6.00e-03 | 6.55e-04 | 676 | -9.16 | 6.29e-19 | -7.28e-03 | -4.71e-03 |
|  | LH | Age x ROI | 1.38e-05 | 3.55e-06 | 676 | 3.87 | 1.18e-04 | 6.78e-06 | 2.07e-05 |
|  | RH | Age | 6.22e-04 | 1.79e-05 | 676 | 34.72 | 2.23e-152 | 5.87e-04 | 6.57e-04 |
|  | RH | ROI | -5.80e-03 | 5.93e-04 | 676 | -9.78 | 3.18e-21 | -6.96e-03 | -4.64e-03 |
|  | RH | Age x ROI | 2.12e-05 | 3.22e-06 | 676 | 6.59 | 8.63e-11 | 1.49e-05 | 2.75e-05 |

**Supplementary Table 3.** Statistical significance of linear mixed models (LMMs), quantifying the relationship between microstructure (R_1_ [s^−1^)) and age [log10 (age in days)) in individual gray matter (GM) regions of interest (ROls) of the dorsal visual stream. LMM: *Rl* ∼ *log10(age of infant) + (1*|*Infant).* Intercept units: s^−1^; Slope units: [s^−1^/log10 (age in days)]. Statistics: *Rsquared* = proportion of variance explained; *OF:* degrees of freedom; *tstats:* t-statistics; *CI* = confidence intervals. *LH/RH:* left/right hemisphere.

| ROI (GM) | Hemi | Intercept | Slope | pValue | Rsquared | DF | tstats | CI lower | CI upper |
| --- | --- | --- | --- | --- | --- | --- | --- | --- | --- |
| V1 | RH | 0.36 | 0.10 | 2.89e-25 | 0.79 | 83 | 14.95 | 8.81e-02 | 1.15e-01 |
| V2 | RH | 0.32 | 0.11 | 6.87e-32 | 0.88 | 83 | 18.95 | 1.00e-01 | 1.24e-01 |
| V3 | RH | 0.30 | 0.12 | 8.22e-38 | 0.92 | 83 | 23.04 | 1.11e-01 | 1.32e-01 |
| V3a | RH | 0.28 | 0.13 | 1.65e-36 | 0.92 | 83 | 22.09 | 1.18e-01 | 1.42e-01 |
| V3b | RH | 0.30 | 0.12 | 9.16e-33 | 0.92 | 83 | 19.52 | 1.08e-01 | 1.33e-01 |
| IPS0 | RH | 0.29 | 0.12 | 8.46e-34 | 0.91 | 83 | 20.21 | 1.10e-01 | 1.33e-01 |
| IPS1 | RH | 0.28 | 0.13 | 1.72e-37 | 0.93 | 83 | 22.80 | 1.17e-01 | 1.39e-01 |
| IPS2 | RH | 0.31 | 0.11 | 1.09e-30 | 0.89 | 83 | 18.18 | 9.36e-02 | 1.17e-01 |
| V1 | LH | 0.35 | 0.10 | 3.47e-27 | 0.82 | 83 | 16.06 | 8.96e-02 | 1.15e-01 |
| V2 | LH | 0.33 | 0.11 | 4.76e-30 | 0.87 | 83 | 17.78 | 9.77e-02 | 1.22e-01 |
| V3 | LH | 0.30 | 0.12 | 6.10e-40 | 0.94 | 83 | 24.65 | 1.13e-01 | 1.33e-01 |
| V3a | LH | 0.28 | 0.13 | 4.87e-38 | 0.94 | 83 | 23.20 | 1.22e-01 | 1.44e-01 |
| V3b | LH | 0.31 | 0.11 | 1.02e-24 | 0.83 | 83 | 14.64 | 9.59e-02 | 1.26e-01 |
| IPS0 | LH | 0.27 | 0.13 | 1.49e-42 | 0.96 | 83 | 26.74 | 1.24e-01 | 1.44e-01 |
| IPS1 | LH | 0.27 | 0.13 | 3.06e-38 | 0.92 | 83 | 23.35 | 1.18e-01 | 1.40e-01 |
| IPS2 | LH | 0.29 | 0.11 | 5.30e-31 | 0.88 | 83 | 18.38 | 9.90e-02 | 1.23e-01 |

**Supplementary Table 4.**
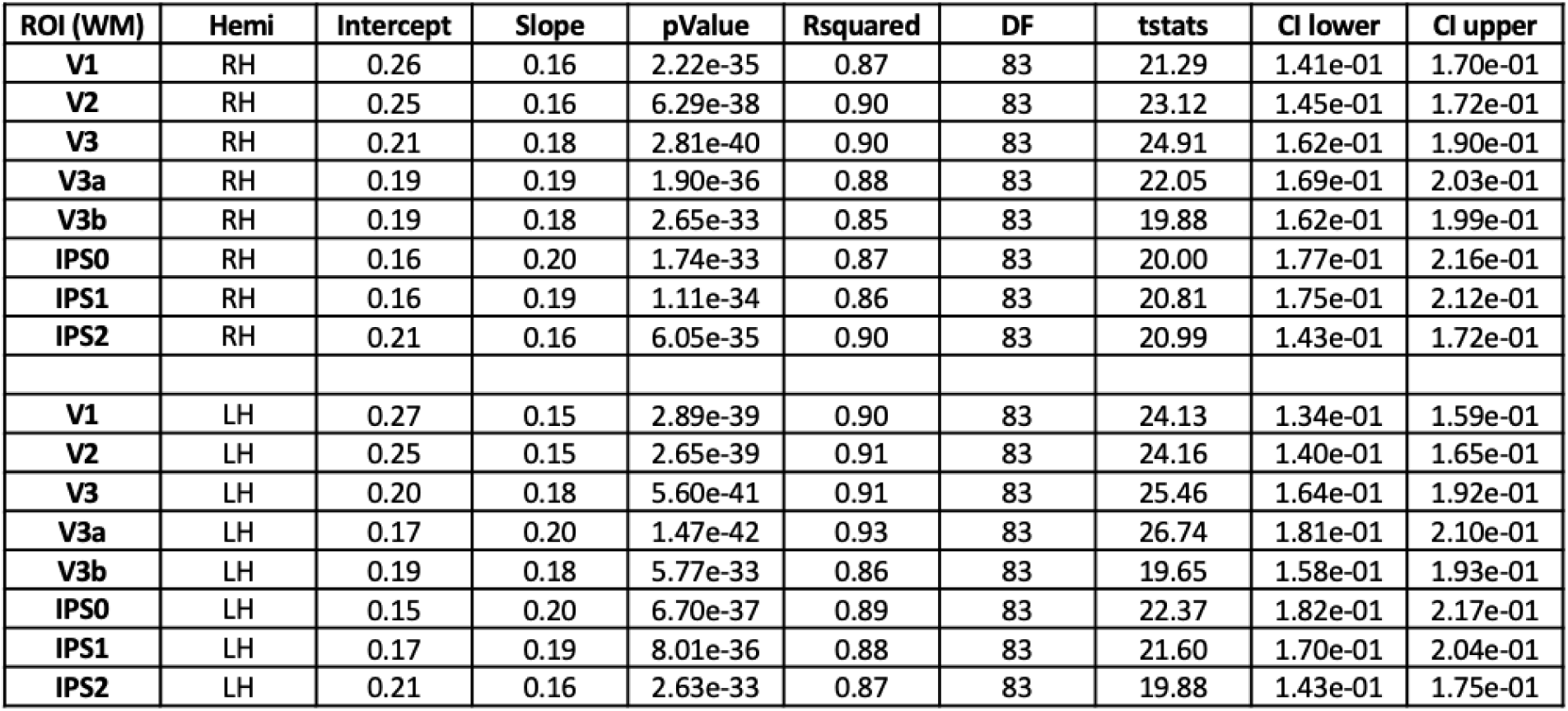
Statistical significance of linear mixed models (LMMs), quantifying the relationship between microstructure (R_1_ [s^−1^]) and age [log10 (age in days)) in individual white matter (WM) regions of interest (ROls) of the dorsal visual stream. LMM: *R1* ∼ *log10(age of infant)+ (1*|*Infant).* Intercept units: s^−1^; Slope units: [s^−1^ /log10 (age in days)]. Statistics: *Rsquared* = proportion of variance explained; *OF:* degrees of freedom; *tstats:* t-statistics; *CI* = confidence intervals. *LH/RH:* left/right hemisphere.

| ROI (WM) | Hemi | Intercept | Slope | pValue | Rsquared | DF | tstats | CI lower | CI upper |
| --- | --- | --- | --- | --- | --- | --- | --- | --- | --- |
| V1 | RH | 0.26 | 0.16 | 2.22e-35 | 0.87 | 83 | 21.29 | 1.41e-01 | 1.70e-01 |
| V2 | RH | 0.25 | 0.16 | 6.29e-38 | 0.90 | 83 | 23.12 | 1.45e-01 | 1.72e-01 |
| V3 | RH | 0.21 | 0.18 | 2.81e-40 | 0.90 | 83 | 24.91 | 1.62e-01 | 1.90e-01 |
| V3a | RH | 0.19 | 0.19 | 1.90e-36 | 0.88 | 83 | 22.05 | 1.69e-01 | 2.03e-01 |
| V3b | RH | 0.19 | 0.18 | 2.65e-33 | 0.85 | 83 | 19.88 | 1.62e-01 | 1.99e-01 |
| IPS0 | RH | 0.16 | 0.20 | 1.74e-33 | 0.87 | 83 | 20.00 | 1.77e-01 | 2.16e-01 |
| IPS1 | RH | 0.16 | 0.19 | 1.11e-34 | 0.86 | 83 | 20.81 | 1.75e-01 | 2.12e-01 |
| IPS2 | RH | 0.21 | 0.16 | 6.05e-35 | 0.90 | 83 | 20.99 | 1.43e-01 | 1.72e-01 |
| V1 | LH | 0.27 | 0.15 | 2.89e-39 | 0.90 | 83 | 24.13 | 1.34e-01 | 1.59e-01 |
| V2 | LH | 0.25 | 0.15 | 2.65e-39 | 0.91 | 83 | 24.16 | 1.40e-01 | 1.65e-01 |
| V3 | LH | 0.20 | 0.18 | 5.60e-41 | 0.91 | 83 | 25.46 | 1.64e-01 | 1.92e-01 |
| V3a | LH | 0.17 | 0.20 | 1.47e-42 | 0.93 | 83 | 26.74 | 1.81e-01 | 2.10e-01 |
| V3b | LH | 0.19 | 0.18 | 5.77e-33 | 0.86 | 83 | 19.65 | 1.58e-01 | 1.93e-01 |
| IPS0 | LH | 0.15 | 0.20 | 6.70e-37 | 0.89 | 83 | 22.37 | 1.82e-01 | 2.17e-01 |
| IPS1 | LH | 0.17 | 0.19 | 8.01e-36 | 0.88 | 83 | 21.60 | 1.70e-01 | 2.04e-01 |
| IPS2 | LH | 0.21 | 0.16 | 2.63e-33 | 0.87 | 83 | 19.88 | 1.43e-01 | 1.75e-01 |

**Supplementary Table 5.**
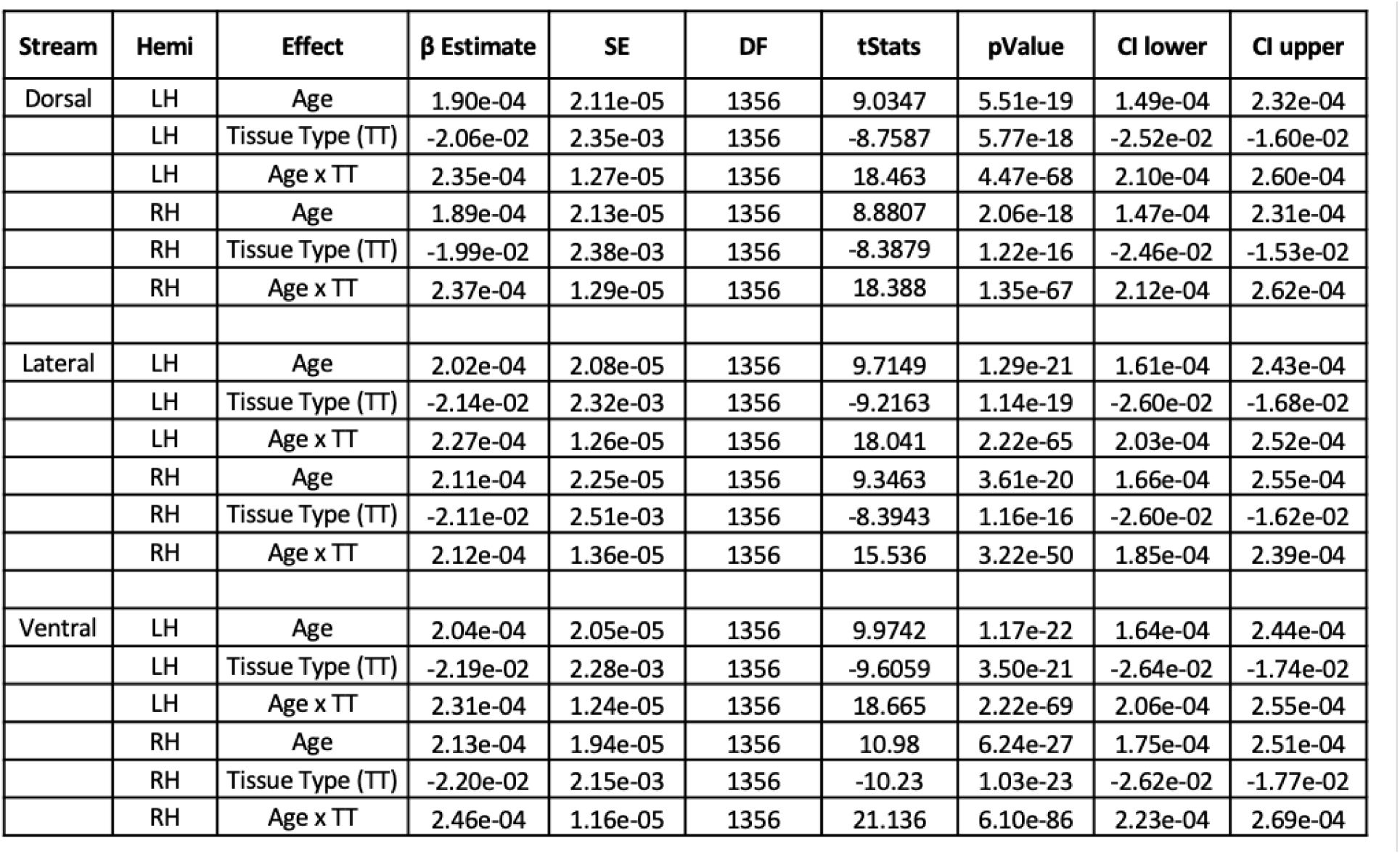
Statistical significance per linear mixed model (LMM). LMMs quantifying the relationship between microstructure (R_1_ [s^−1^]) and age [log10 (age in days)] and tissue type (TT) (gray matter(GM)/white matter (WM)). LMM: *R1* ∼ *log10(age of infant) × TT+ (1*|*Infant).* Three effects are shown per model per hemisphere: main effects of age and TT, and interaction between age and TT. Statistics: *SE:* Standard error; *OF:* degrees of freedom; *tstats:* t-statistics; *CI* = confidence intervals. *LH/RH:* left/right hemisphere.

| Stream | Hemi | Effect | $\beta$ Estimate | SE | DF | tStats | pValue | CI lower | CI upper |
| --- | --- | --- | --- | --- | --- | --- | --- | --- | --- |
| Dorsal | LH | Age | 1.90e-04 | 2.11e-05 | 1356 | 9.0347 | 5.51e-19 | 1.49e-04 | 2.32e-04 |
|  | LH | Tissue Type (TT) | -2.06e-02 | 2.35e-03 | 1356 | -8.7587 | 5.77e-18 | -2.52e-02 | -1.60e-02 |
|  | LH | Age x TT | 2.35e-04 | 1.27e-05 | 1356 | 18.463 | 4.47e-68 | 2.10e-04 | 2.60e-04 |
|  | RH | Age | 1.89e-04 | 2.13e-05 | 1356 | 8.8807 | 2.06e-18 | 1.47e-04 | 2.31e-04 |
|  | RH | Tissue Type (TT) | -1.99e-02 | 2.38e-03 | 1356 | -8.3879 | 1.22e-16 | -2.46e-02 | -1.53e-02 |
|  | RH | Age x TT | 2.37e-04 | 1.29e-05 | 1356 | 18.388 | 1.35e-67 | 2.12e-04 | 2.62e-04 |
| Lateral | LH | Age | 2.02e-04 | 2.08e-05 | 1356 | 9.7149 | 1.29e-21 | 1.61e-04 | 2.43e-04 |
|  | LH | Tissue Type (TT) | -2.14e-02 | 2.32e-03 | 1356 | -9.2163 | 1.14e-19 | -2.60e-02 | -1.68e-02 |
|  | LH | Age x TT | 2.27e-04 | 1.26e-05 | 1356 | 18.041 | 2.22e-65 | 2.03e-04 | 2.52e-04 |
|  | RH | Age | 2.11e-04 | 2.25e-05 | 1356 | 9.3463 | 3.61e-20 | 1.66e-04 | 2.55e-04 |
|  | RH | Tissue Type (TT) | -2.11e-02 | 2.51e-03 | 1356 | -8.3943 | 1.16e-16 | -2.60e-02 | -1.62e-02 |
|  | RH | Age x TT | 2.12e-04 | 1.36e-05 | 1356 | 15.536 | 3.22e-50 | 1.85e-04 | 2.39e-04 |
| Ventral | LH | Age | 2.04e-04 | 2.05e-05 | 1356 | 9.9742 | 1.17e-22 | 1.64e-04 | 2.44e-04 |
|  | LH | Tissue Type (TT) | -2.19e-02 | 2.28e-03 | 1356 | -9.6059 | 3.50e-21 | -2.64e-02 | -1.74e-02 |
|  | LH | Age x TT | 2.31e-04 | 1.24e-05 | 1356 | 18.665 | 2.22e-69 | 2.06e-04 | 2.55e-04 |
|  | RH | Age | 2.13e-04 | 1.94e-05 | 1356 | 10.98 | 6.24e-27 | 1.75e-04 | 2.51e-04 |
|  | RH | Tissue Type (TT) | -2.20e-02 | 2.15e-03 | 1356 | -10.23 | 1.03e-23 | -2.62e-02 | -1.77e-02 |
|  | RH | Age x TT | 2.46e-04 | 1.16e-05 | 1356 | 21.136 | 6.10e-86 | 2.23e-04 | 2.69e-04 |

**Supplementary Table 6.**
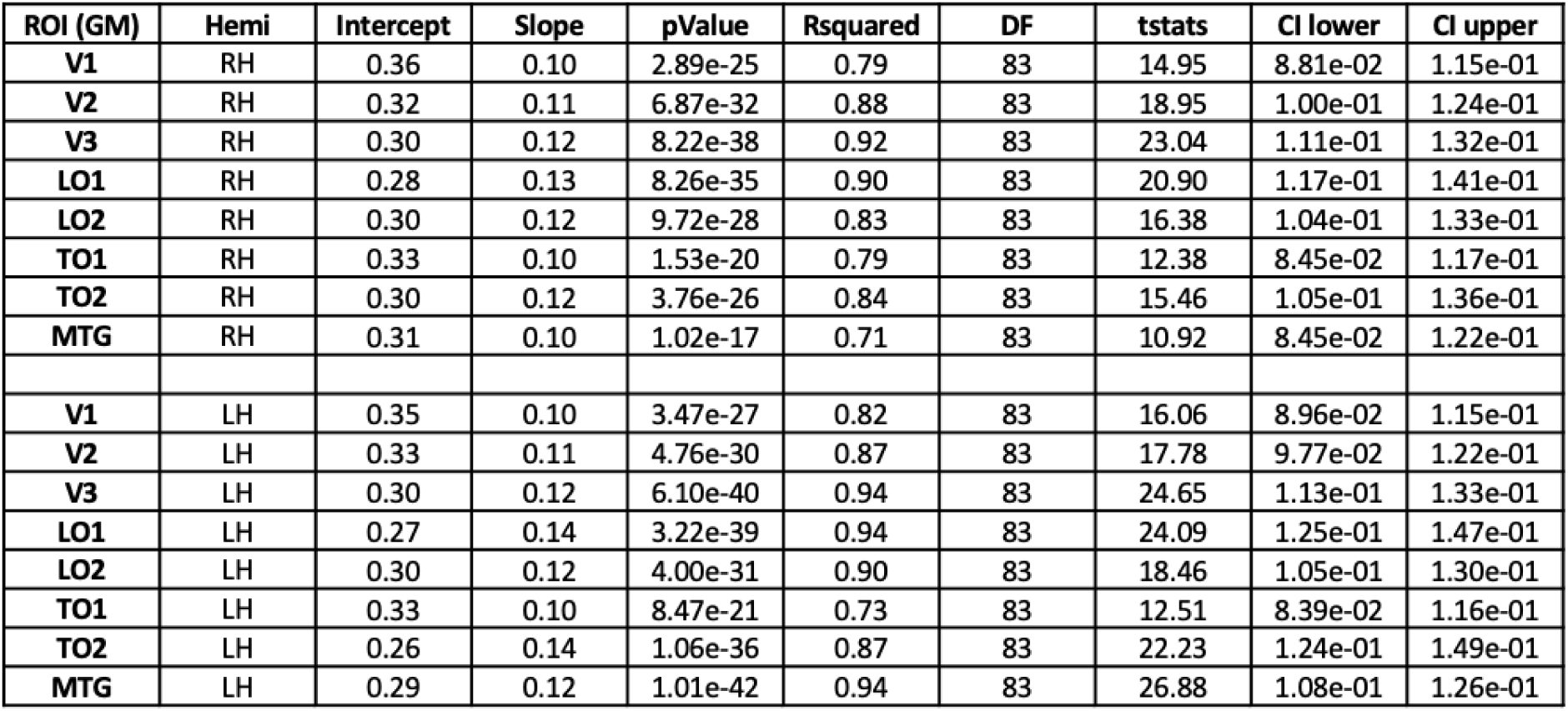
Statistical significance of linear mixed models (LMMs), quantifying the relationship between microstructure (R_1_ [s^−1^]) and age [log10 (age in days)] in individual gray matter (GM) regions of interest (ROls) of the lateral visual stream. LMM: *R1* ∼ *log10(age of infant)+ (1*|*Infant).* Intercept units: s^−1^; Slope units: [s^−1^ /log10 (age in days)]. Statistics: *Rsquared* = proportion of variance explained; *OF:* degrees of freedom; *tstats:* t-statistics; *CI* = confidence intervals. *LH/RH:* left/right hemisphere.

**Supplementary Table 7.** Statistical significance of linear mixed models (LMMs), quantifying the relationship between microstructure {R_1_ [s^−1^]) and age [log10 (age in days)] in individual white matter **(WM)** regions of interest (ROls) of the lateral visual stream. LMM: *R1* ∼ *log10(age of infant)+ (1*|*Infant).* Intercept units: s^−1^; Slope units: [s^−1^ /log10 (age in days)]. Statistics: *Rsquared* = proportion of variance explained; *OF:* degrees of freedom; *tstats:* t-statistics; *CI=* confidence intervals. *LH/RH:* left/right hemisphere.

| ROI (WM) | Hemi | Intercept | Slope | pValue | Rsquared | DF | tstats | CI lower | CI upper |
| --- | --- | --- | --- | --- | --- | --- | --- | --- | --- |
| V1 | RH | 0.26 | 0.16 | 2.22e-35 | 0.87 | 83 | 21.29 | 1.41e-01 | 1.70e-01 |
| V2 | RH | 0.25 | 0.16 | 6.29e-38 | 0.90 | 83 | 23.12 | 1.45e-01 | 1.72e-01 |
| V3 | RH | 0.21 | 0.18 | 2.81e-40 | 0.90 | 83 | 24.91 | 1.62e-01 | 1.90e-01 |
| LO1 | RH | 0.17 | 0.19 | 4.03e-37 | 0.88 | 83 | 22.53 | 1.78e-01 | 2.12e-01 |
| LO2 | RH | 0.20 | 0.17 | 4.38e-36 | 0.88 | 83 | 21.79 | 1.56e-01 | 1.88e-01 |
| TO1 | RH | 0.23 | 0.15 | 8.96e-33 | 0.85 | 83 | 19.53 | 1.37e-01 | 1.68e-01 |
| TO2 | RH | 0.18 | 0.19 | 1.59e-28 | 0.81 | 83 | 16.85 | 1.63e-01 | 2.07e-01 |
| MTG | RH | 0.23 | 0.14 | 8.24e-27 | 0.81 | 83 | 15.84 | 1.25e-01 | 1.61e-01 |
| V1 | LH | 0.27 | 0.15 | 2.89e-39 | 0.90 | 83 | 24.13 | 1.34e-01 | 1.59e-01 |
| V2 | LH | 0.25 | 0.15 | 2.65e-39 | 0.91 | 83 | 24.16 | 1.40e-01 | 1.65e-01 |
| V3 | LH | 0.20 | 0.18 | 5.60e-41 | 0.91 | 83 | 25.46 | 1.64e-01 | 1.92e-01 |
| LO1 | LH | 0.15 | 0.20 | 1.70e-35 | 0.87 | 83 | 21.37 | 1.84e-01 | 2.22e-01 |
| LO2 | LH | 0.19 | 0.17 | 1.73e-36 | 0.89 | 83 | 22.07 | 1.59e-01 | 1.90e-01 |
| TO1 | LH | 0.20 | 0.17 | 3.48e-36 | 0.85 | 83 | 21.86 | 1.53e-01 | 1.83e-01 |
| TO2 | LH | 0.16 | 0.19 | 1.11e-32 | 0.81 | 83 | 19.47 | 1.73e-01 | 2.12e-01 |
| MTG | LH | 0.19 | 0.17 | 3.30e-34 | 0.86 | 83 | 20.49 | 1.52e-01 | 1.85e-01 |

**Supplementary Table 8.** Statistical significance of linear mixed models (LMMs), quantifying the relationship between microstructure (R_1_ [s^−1^]) and age [logl0 (age in days)] in individual gray matter (GM) regions of interest (ROls) of the ventral visual stream. LMM: *R1* ∼ *log10(age of infant) +(1*|*Infant).* Intercept units: s^−1^; Slope units: [s^−1^ /log10 (age in days)]. Statistics: *Rsquared* = proportion of variance explained; *DF:* degrees of freedom; *tstats:* t-statistics; *CI=* confidence intervals. *LH/RH:* left/right hemisphere.

| ROI (GM) | Hemi | Intercept | Slope | pValue | Rsquared | DF | tstats | CI lower | CI upper |
| --- | --- | --- | --- | --- | --- | --- | --- | --- | --- |
| V1 | RH | 0.36 | 0.10 | 2.89e-25 | 0.79 | 83 | 14.95 | 8.81e-02 | 1.15e-01 |
| V2 | RH | 0.32 | 0.11 | 6.87e-32 | 0.88 | 83 | 18.95 | 1.00e-01 | 1.24e-01 |
| V3 | RH | 0.30 | 0.12 | 8.22e-38 | 0.92 | 83 | 23.04 | 1.11e-01 | 1.32e-01 |
| hV4 | RH | 0.29 | 0.12 | 1.76e-26 | 0.76 | 83 | 15.65 | 1.09e-01 | 1.41e-01 |
| V01 | RH | 0.28 | 0.13 | 2.73e-33 | 0.86 | 83 | 19.87 | 1.15e-01 | 1.41e-01 |
| V02 | RH | 0.29 | 0.12 | 1.55e-35 | 0.87 | 83 | 21.40 | 1.13e-01 | 1.36e-01 |
| PHC1 | RH | 0.30 | 0.12 | 4.43e-34 | 0.86 | 83 | 20.40 | 1.09e-01 | 1.32e-01 |
| PHC2 | RH | 0.29 | 0.12 | 1.75e-30 | 0.82 | 83 | 18.06 | 1.07e-01 | 1.34e-01 |
| V1 | LH | 0.35 | 0.10 | 3.47e-27 | 0.82 | 83 | 16.06 | 8.96e-02 | 1.15e-01 |
| V2 | LH | 0.33 | 0.11 | 4.76e-30 | 0.87 | 83 | 17.78 | 9.77e-02 | 1.22e-01 |
| V3 | LH | 0.30 | 0.12 | 6.10e-40 | 0.94 | 83 | 24.65 | 1.13e-01 | 1.33e-01 |
| hV4 | LH | 0.27 | 0.13 | 1.08e-32 | 0.86 | 83 | 19.47 | 1.21e-01 | 1.48e-01 |
| V01 | LH | 0.28 | 0.13 | 1.03e-37 | 0.92 | 83 | 22.96 | 1.18e-01 | 1.40e-01 |
| V02 | LH | 0.29 | 0.12 | 1.05e-40 | 0.94 | 83 | 25.24 | 1.11e-01 | 1.30e-01 |
| PHC1 | LH | 0.29 | 0.12 | 6.75e-39 | 0.92 | 83 | 23.85 | 1.12e-01 | 1.32e-01 |
| PHC2 | LH | 0.30 | 0.12 | 2.92e-32 | 0.85 | 83 | 19.19 | 1.04e-01 | 1.28e-01 |

**Supplementary Table 9.**
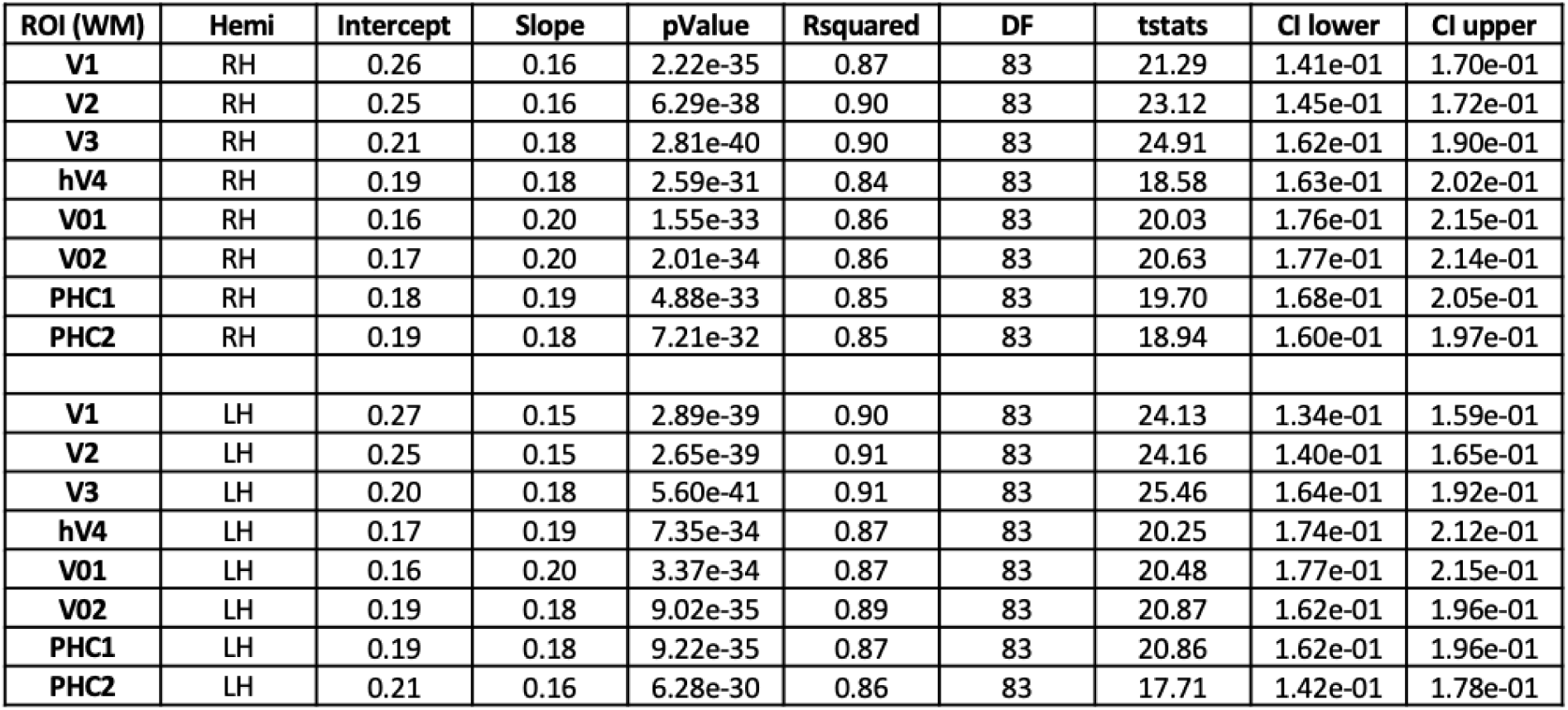
Statistical significance of linear mixed models (LMMs), quantifying the relationship between microstructure (R_1_ [s^−1^]) and age [logl0 (age in days)] in individual white matter (WM) regions of interest (ROls) of the ventral visual stream. LMM: *R1* ∼ *log10(age of infant)+ (1*|*Infant).* Intercept units: s^−1^; Slope units: [s^−1^ /log10 (age in days)). Statistics: *Rsquared* = proportion of variance explained; *DF:* degrees of freedom; *tstats:* t-statistics; *CI* = confidence intervals. *LH/RH:* left/right hemisphere.

| ROI (WM) | Hemi | Intercept | Slope | pValue | Rsquared | DF | tstats | CI lower | CI upper |
| --- | --- | --- | --- | --- | --- | --- | --- | --- | --- |
| V1 | RH | 0.26 | 0.16 | 2.22e-35 | 0.87 | 83 | 21.29 | 1.41e-01 | 1.70e-01 |
| V2 | RH | 0.25 | 0.16 | 6.29e-38 | 0.90 | 83 | 23.12 | 1.45e-01 | 1.72e-01 |
| V3 | RH | 0.21 | 0.18 | 2.81e-40 | 0.90 | 83 | 24.91 | 1.62e-01 | 1.90e-01 |
| hV4 | RH | 0.19 | 0.18 | 2.59e-31 | 0.84 | 83 | 18.58 | 1.63e-01 | 2.02e-01 |
| V01 | RH | 0.16 | 0.20 | 1.55e-33 | 0.86 | 83 | 20.03 | 1.76e-01 | 2.15e-01 |
| V02 | RH | 0.17 | 0.20 | 2.01e-34 | 0.86 | 83 | 20.63 | 1.77e-01 | 2.14e-01 |
| PHC1 | RH | 0.18 | 0.19 | 4.88e-33 | 0.85 | 83 | 19.70 | 1.68e-01 | 2.05e-01 |
| PHC2 | RH | 0.19 | 0.18 | 7.21e-32 | 0.85 | 83 | 18.94 | 1.60e-01 | 1.97e-01 |
| V1 | LH | 0.27 | 0.15 | 2.89e-39 | 0.90 | 83 | 24.13 | 1.34e-01 | 1.59e-01 |
| V2 | LH | 0.25 | 0.15 | 2.65e-39 | 0.91 | 83 | 24.16 | 1.40e-01 | 1.65e-01 |
| V3 | LH | 0.20 | 0.18 | 5.60e-41 | 0.91 | 83 | 25.46 | 1.64e-01 | 1.92e-01 |
| hV4 | LH | 0.17 | 0.19 | 7.35e-34 | 0.87 | 83 | 20.25 | 1.74e-01 | 2.12e-01 |
| V01 | LH | 0.16 | 0.20 | 3.37e-34 | 0.87 | 83 | 20.48 | 1.77e-01 | 2.15e-01 |
| V02 | LH | 0.19 | 0.18 | 9.02e-35 | 0.89 | 83 | 20.87 | 1.62e-01 | 1.96e-01 |
| PHC1 | LH | 0.19 | 0.18 | 9.22e-35 | 0.87 | 83 | 20.86 | 1.62e-01 | 1.96e-01 |
| PHC2 | LH | 0.21 | 0.16 | 6.28e-30 | 0.86 | 83 | 17.71 | 1.42e-01 | 1.78e-01 |

## Notes

### Competing Interest Statement

The authors have declared no competing interest.

